# Dynamics of *hunchback* translation in real time and at single mRNA resolution in the *Drosophila* embryo

**DOI:** 10.1101/2021.02.16.430474

**Authors:** Daisy J. Vinter, Caroline Hoppe, Thomas G. Minchington, Catherine Sutcliffe, Hilary L. Ashe

**Author notes:** Institute of Science and Technology Austria (IST Austria), Am Campus 1, 3400 Klosterneuburg, Austria.

## Abstract

The Hunchback (Hb) transcription factor is critical for anterior-posterior patterning of the *Drosophila* embryo. Despite the maternal *hb* mRNA acting as a paradigm for translational regulation, due to its repression in the posterior of the embryo, little is known about the translatability of zygotically transcribed *hb* mRNAs. Here we adapt the SunTag system, developed for imaging translation at single mRNA resolution in tissue culture cells, to the *Drosophila* embryo to study the translation dynamics of zygotic *hb* mRNAs. Using singlemolecule imaging in fixed and live embryos, we provide evidence for translational repression of zygotic *SunTag-hb* mRNAs. While the proportion of *SunTag-hb* mRNAs translated is initially uniform, translation declines from the anterior over time until it becomes restricted to a posterior band in the expression domain. We discuss how regulated *hb* mRNA translation may help establish the sharp Hb expression boundary, which is a model for precision and noise during developmental patterning. Overall, our data show how use of the SunTag method on fixed and live embryos is a powerful combination for elucidating spatiotemporal regulation of mRNA translation in *Drosophila*.

## Introduction

The gap gene *hunchback (hb)* is essential for body plan specification during *Drosophila* embryogenesis, primarily by establishing the gene expression boundaries of other gap genes through transcriptional repression (Hulskamp et al., 1990; Struhl et al., 1992). Embryos with complete loss of *hb* function lack all gnathal and thoracic segments, show reversed polarity of the first 2-3 abdominal segments and the seventh and eighth abdominal segments are fused (Lehmann and Nusslein-Volhard, 1987). *hb* is maternally and zygotically expressed (Bender et al., 1988; Tautz et al., 1987). Maternally expressed *hb* mRNA is uniformly distributed in the embryo, but its translation is repressed in the posterior by Nanos, Pumilio, Brain Tumor and d4EHP (Cho et al., 2006; Hülskamp et al., 1989; Irish et al., 1989; Murata and Wharton, 1995; Sonoda and Wharton, 1999, 2001; Struhl, 1989; Wharton and Struhl, 1991).

Zygotic *hb* expression is dynamic with spatial and temporal regulation relying on two promoters and three enhancers. Initially *hb* is transcribed from the proximal P2 promoter (Margolis et al.; Schröder et al., 1988) in a broad anterior domain in nuclear cycle (nc) 11 embryos (Bender et al., 1988; Tautz et al., 1987; Tautz and Pfeifle, 1989), in response to the Bicoid (Bcd) gradient (Driever and Nüsslein-Volhard, 1989; Struhl et al., 1989). This expression in the anterior half of the embryo persists into early nc14, when *hb* mRNAs from the P2 promoter also appear transiently in a posterior cap. From mid-nc14, both the P1 and P2 promoters are active with the resulting transcripts accumulating in a stripe positioned at the posterior boundary of the anterior domain of *hb* expression, around 50% egg length (EL), as well as in the posterior region of the embryo (Margolis et al., 1995). By this time, the broad anterior domain of *hb* mRNA has decayed so that only two weaker stripes of mRNA are detectable in the anterior, in addition to the newly activated central stripe (Margolis et al., 1995; Tautz and Pfeifle, 1989). The early *hb* expression in the anterior half of the embryo is controlled by the Bcd-responsive proximal and shadow enhancers (Driever and Nüsslein-Volhard, 1989; Margolis et al., 1995; Perry et al., 2011; Struhl et al., 1989), whereas a stripe enhancer activates transcription in both the central and posterior stripes (Margolis et al., 1995; Perry et al., 2012). The stripe enhancer is ubiquitously activated, but two stripes are formed due to the action of repressors encoded by other gap genes (Perry et al., 2012).

Hb protein distribution mirrors that of the zygotic mRNA pattern described above, except that the protein persists in the entire anterior half of mid-late nc14 embryos, likely due to slower degradation of the protein than mRNA (Perry et al., 2012; Tautz and Pfeifle, 1989; Wu et al., 2001). The anterior Hb domain has a steep boundary of Hb protein in the centre of the embryo, in part due to expression from the stripe enhancer (Perry et al., 2012). Overall, much is known about the transcriptional regulation of *hb*, including quantitative live imaging studies of the output from the proximal enhancer and P2 promoter, with nascent transcription detected as early as nc 9-10 (Garcia et al., 2013; Lucas et al., 2013). In comparison, little is known about the translatability of zygotic *hb* mRNAs in space and time during early embryogenesis.

The *Drosophila* embryo represents a paradigm for understanding how translational control can underpin developmental patterning, for example through studies of the translational repression of the *oskar, caudal* and maternal *hb* mRNAs (Lasko, 2012). In addition, ribosome profiling has provided new insights into the control of translation of the entire transcriptome in *Drosophila* embryos (Dunn et al., 2013; Eichhorn et al., 2016). However, the temporal dynamics of mRNA translation have been less well studied due to the absence of a method to image and quantitate translation at a single mRNA level in *Drosophila*. Recently the SunTag methodology has been developed in tissue culture cells, in which 24 copies of a GCN4 peptide, the SunTag, is introduced at the start of the coding sequence. This SunTag array can be detected as it emerges from the ribosome exit channel, by the binding of a co-expressed single-chain variable fragment antibody (scFv) fused to GFP (Pichon et al., 2016; Tanenbaum et al., 2014; Wang et al., 2016; Wu et al., 2016; Yan et al., 2016). In combination with the MS2 system to visualise individual mRNAs, translation can be followed in real time and quantified. As a result, this method has provided powerful new insights into the regulation of translation; for example, localisation or isoform dependent differences in translation efficiency (Voigt et al., 2017; Wu et al., 2016; Yan et al., 2016).

By adapting the SunTag system for *Drosophila* embryos, we have studied the translation dynamics of zygotic *hb* mRNAs in both fixed and live embryos, which reveals temporal regulation of *hb* mRNA translation in the early embryo. Translation is uniform across the *hb* expression domain in nc12-13, but then becomes repressed in the anterior so that *hb* mRNA translation persists in only a band of cells at the posterior of the expression domain in nc14. These results suggest a previously unknown translational regulation of zygotic *hb* mRNAs, which may contribute to the refinement and precision of the Hb protein boundary, impacting downstream developmental patterning.

## Results

In order to visualise translation of single mRNAs in the early *Drosophila* embryo we focused on the gap gene *hb* and adapted the SunTag method that has previously been used to monitor translation of mRNAs in tissue culture cells and cultured neurons (Pichon et al., 2016; Wang et al., 2016; Wu et al., 2016; Yan et al., 2016). We generated a *SunTag-hb* reporter gene, under the control of the P2 promoter and primary Bicoid-responsive enhancer (Driever and Nüsslein-Volhard, 1989; Struhl et al., 1989), which carries an array of 24 copies of the SunTag peptide at its N terminus (Fig. 1A). This peptide is recognised by a single-chain antibody (scFv) fused to mNeonGreen (NG), which is expressed as a second transgene, under the control of the *nanos* (*nos*) promoter to allow maternal expression (Fig. 1A). We added a nuclear localization signal to the scFv-NG protein to reduce cytoplasmic accumulation of unbound scFv-NG, as described previously (Pichon et al., 2016), to improve the signal to noise in the cytoplasm and facilitate detection of translation of individual mRNAs. As a result, NG antibody staining of embryos carrying only the scFv-NG transgene shows uniform nuclear NG signal throughout the embryo during early stages (Fig. 1B, C).

**Fig. 1.**
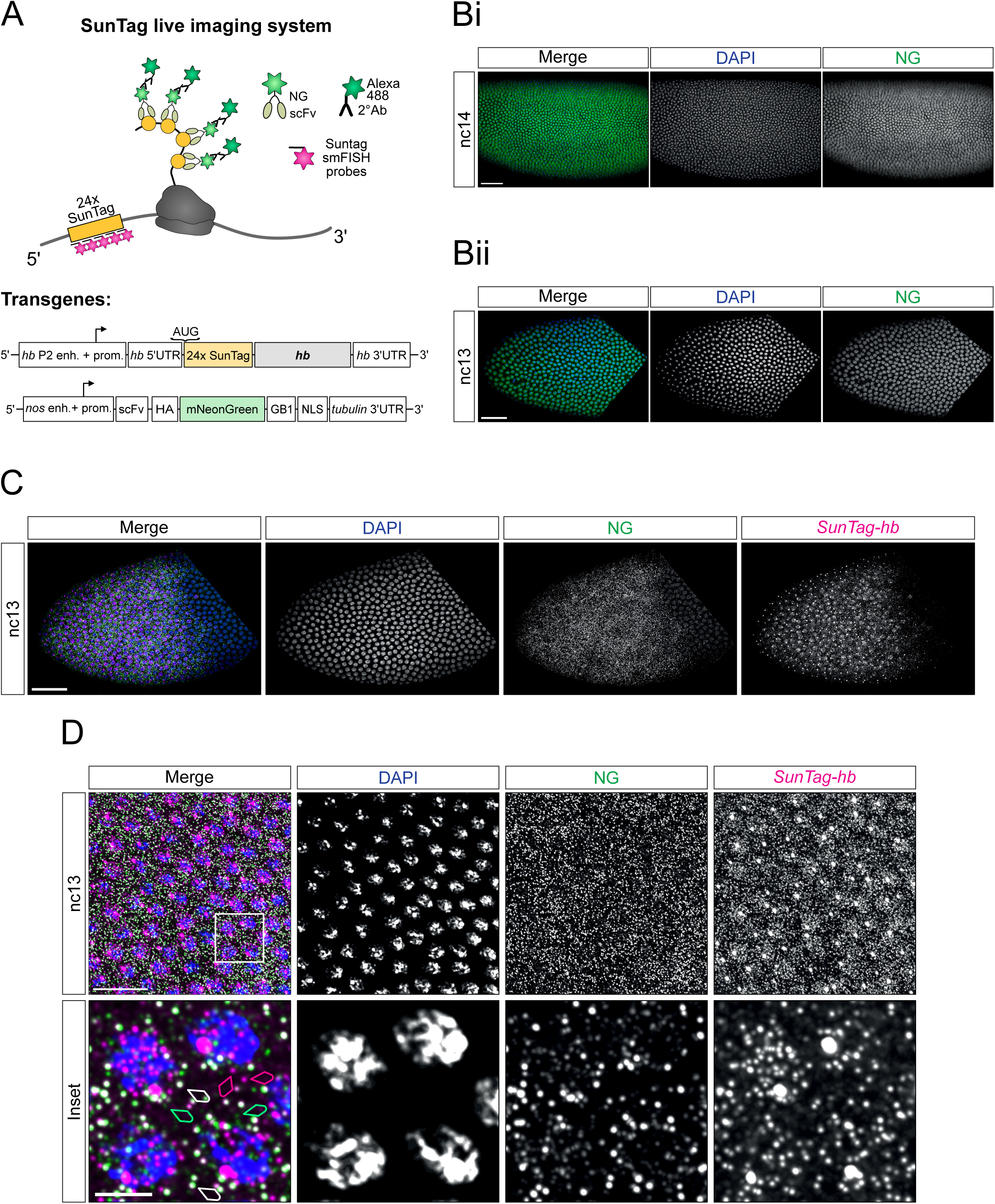
Visualisation of translation of single *SunTag-hb* mRNAs in the *Drosophila* embryo. (A) Schematic showing translation of the *SunTag-hb* mRNA. SunTag peptides are recognised by a scFv-NG fusion protein that is detected by an antibody stain. mRNAs are recognised by smFISH probes against the SunTag sequence. Schematics show the organisation of the transgenes used for visualisation of translation. (B) (Bi) *Drosophila* nc14 embryo from females maternally expressing *scFv-NG* crossed to wild type males, stained with DAPI (blue) and anti-NG (green). Scale bar: 50μm. (Bii) As in (Bi), except staining in the anterior of nc13 embryo is shown. Scale bar: 50μm. (C) Anterior region of a nc13 embryo from females maternally expressing *scFv-NG* crossed to males carrying the *SunTag-hb* transgene, stained with DAPI (blue), anti-NG (green) and *SunTag* smFISH probes (magenta). Scale bar: 50μm. (D) As in (D) but higher magnification views are shown. The bottom panel shows the boxed region of interest, image is a maximum projection of 9 slices. White arrows show colocalised scFv-NG and *SunTag-hb* smFISH spots representing translated mRNAs. Magenta arrows identify uncolocalised *SunTag-hb* smFISH spots, which are untranslated mRNAs. Green arrows show uncolocalised weaker scFv-NG spots that are proteins. Scale bars: 20μm (top panel), 5μm (lower panel).

First, we studied fixed embryos maternally expressing *scFv-NG* and carrying a single copy of the *SunTag-hb* transgene. We used smFISH probes against the SunTag sequences to visualise *SunTag-hb* mRNAs, combined with NG antibody staining to detect the scFv-NG protein (Fig. 1A). Visualisation of *SunTag-hb* mRNAs in a nc13 embryo reveals their localisation in a broad anterior domain, with bright transcription foci and the dimmer signal associated with single mRNAs (Fig. 1D). It is also clear from the scFv-NG staining that, while this protein is predominantly nuclear in the posterior region of the embryo as expected due to the presence of the NLS, cytoplasmic NG signal is observed in the anterior (Fig. 1D, cf Fig. 1C). A higher magnification image from the anterior region of a nc13 embryo allows individual mRNAs to be visualised (Fig. 1E, pink arrowheads). A subset of these mRNAs has a colocalizing NG signal (Fig. 1E, white arrowheads), consistent with these mRNAs being translated. In addition, weaker green puncta are detected that are not colocalised with a mRNA signal, which are likely single proteins within the cytoplasm (Fig. 1E, green arrowheads).

As we can detect translation of single mRNAs, we determined the *hb* mRNA translation efficiency depending on position in the expression domain. Firstly, we quantitated the total number of *SunTag-hb* mRNAs in early nc12 (Fig. 2A). As the embryo is not cellularised at this stage, we assigned mRNAs to the nearest nucleus giving us the mRNA number per nuclear territory. To focus only on cytoplasmic mRNAs and proteins, we masked nuclei to exclude signals in the nucleus from our quantitation (see Methods for more details). The cytoplasmic *SunTag-hb* mRNA number, plotted as the mean from 3 embryos, shows a graded distribution with an increase towards the central region of the *hb* expression domain, followed by a decrease in the posterior half (Fig. 2Bi). This mRNA profile is similar to that described for the mRNA output from the primary *hb* enhancer (Garcia et al., 2013) and for endogenous *hb* mRNAs at nc12 (Little et al., 2013). Quantitation of the number of *SunTag-hb* mRNAs being translated shows a similar graded profile to that of the total mRNA (Fig. 2Bii). Consistent with this, the proportion of *SunTag-hb* mRNAs being translated is constant at ~55% regardless of position within the expression domain (Fig. 2Biii). The mean number of cytoplasmic *SunTag-hb* mRNAs, with the number and proportion translated for each embryo is shown in Fig. S1A, all the data points for a representative embryo are shown in Fig. S1B.

**Figure 2:**
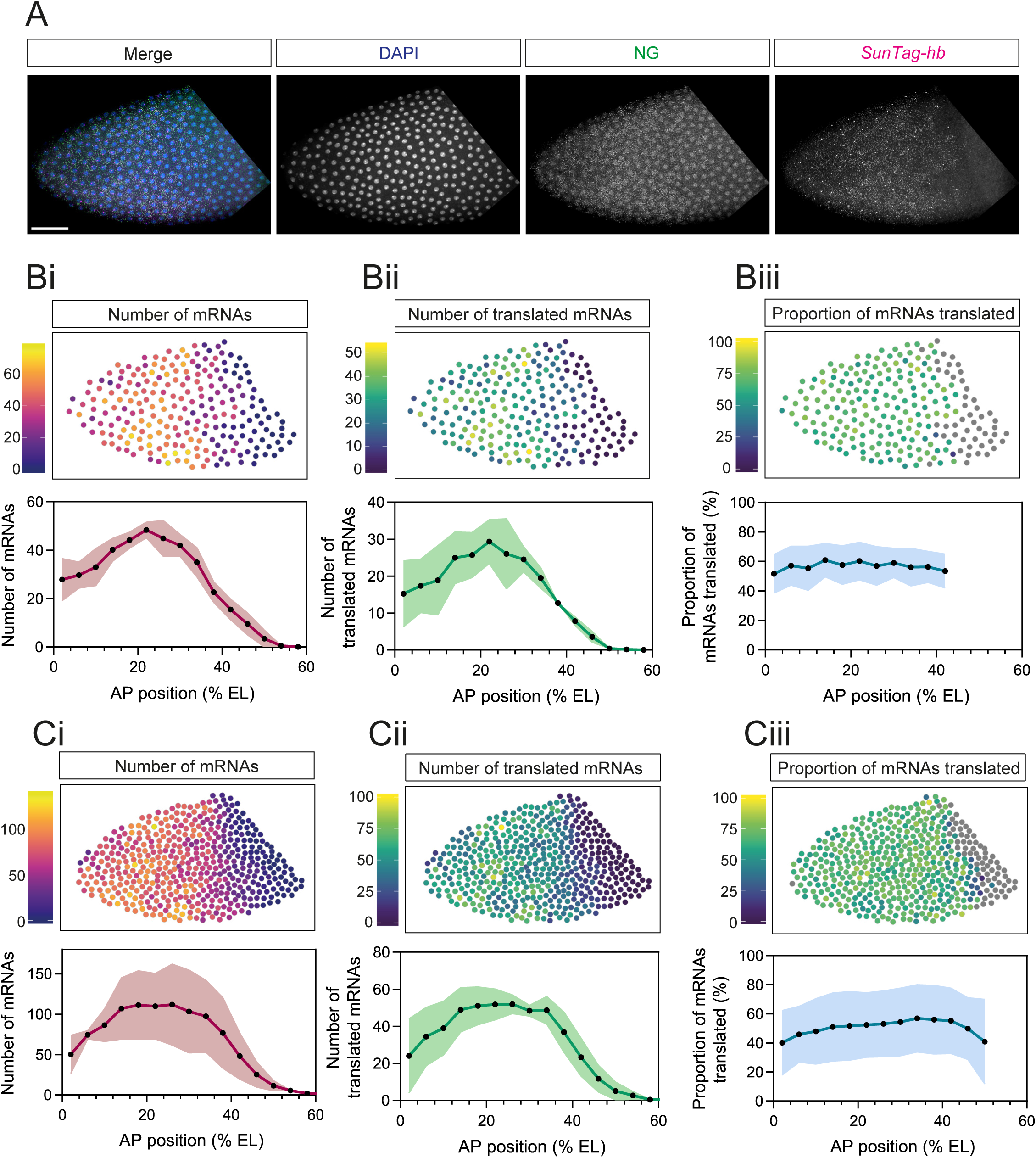
Uniform translation of *SunTag-hb* mRNAs across the expression domain in nc12 and 13. (A) Anterior region of a fixed *Drosophila* nc12 embryo with maternal and zygotic expression of the *scFv-NG* and *SunTag-hb* transgenes, respectively, stained with DAPI (blue), anti-NG (green) and *SunTag* smFISH probes (magenta). Scale bar: 50μm. (B) Quantitation per nuclear territory of total cytoplasmic mRNAs (i), translated mRNAs (ii), and percentage of mRNAs translated (iii) in nc12 embryos. Heat maps correspond to the embryo image in Fig.2A. % EL is estimated based on an embryo length of 500μm. Nuclear territories are binned along the AP axis (bin size 20μm) and mean values per bin are reported. Mean ± SD, n = 3 embryos. (C) As in (B), except the quantitation is for nc13 embryos and the heatmaps are for the embryo shown in Fig. 1D.

To investigate translation of *SunTag-hb* mRNAs through development, we repeated this imaging and quantitation for nc13 embryos (Fig. 2C, the representative heatmap shown is for the embryo image in Fig. 1D). At this stage, the mRNA distribution is similar to that in nc12 although mRNA number has increased (Fig. 2Ci). The numbers of translated mRNAs are also increased and again their distribution mirrors that of the total mRNA profile (Fig. 2Cii), so that the percentage translated is uniform (~50%) across the expression domain (Fig. 2Ciii). The mean numbers of total and translated *SunTag-hb* mRNAs is shown for individual nc13 embryos in Fig. S1C, in addition to the data for each nuclear territory for a representative embryo (Fig. S1D).

Next, we focused on early nc14 embryos, based on the presence of *SunTag-hb* transcription sites, as *hb* is only transcribed from the proximal enhancer early in nc14 (Garcia et al., 2013). Visualisation of *SunTag-hb* mRNAs and NG signal in early nc14 embryos reveals a different distribution of translation sites, with the highest NG signal detected in a posterior band within the expression domain (Fig. 3A). The higher magnification images show the paucity of translation sites in the anterior, despite high mRNA density, compared to the posterior (Fig. 3A). Quantitation of the total number of cytoplasmic *SunTag-hb* mRNAs per nuclear territory in early nc14 (Fig. 3Bi, Fig. S1E-Fi) reveals a similar profile to that observed in nc13, with a small increase in the total mRNA number, again consistent with published data for the output from the *hb* primary enhancer (Bothma et al., 2015; Garcia et al., 2013) and endogenous *hb* at this stage (Little et al., 2013). However, at this stage, the profile of *SunTag-hb* translated mRNAs differs from the mRNA distribution and instead peaks at the posterior of the expression domain (Fig. 3Bii, Fig. S1E-Fii). As such, the proportion of mRNAs translated is less than 10% in the anterior of the expression domain, increasing to ~60% in the posterior (Fig. 3Biii, Fig. S1Eiii). These data suggest that there is an active translation repression mechanism in the anterior at nc14, but not in the posterior where translation of *SunTag-hb* mRNAs persists.

**Figure 3:**
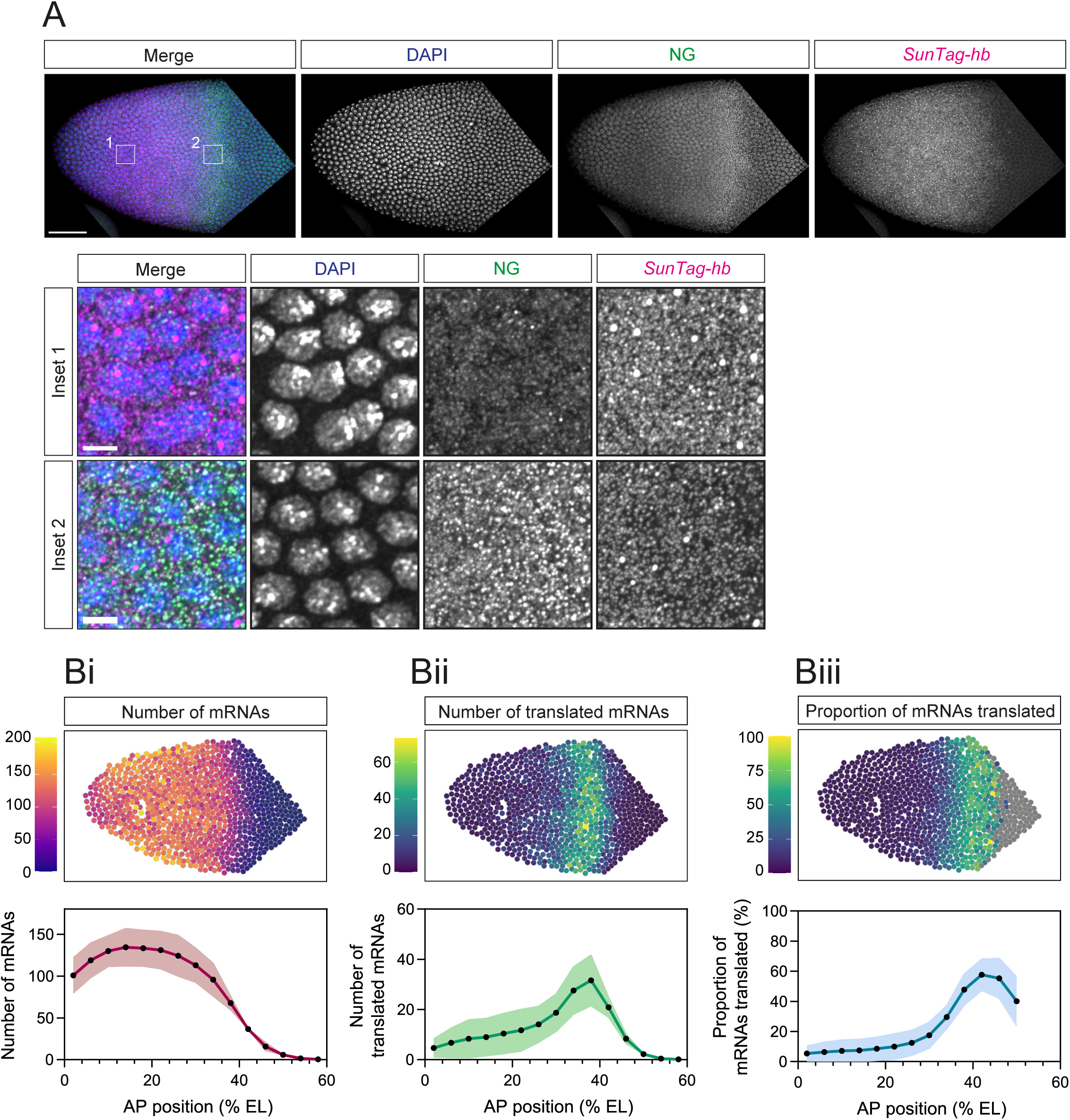
Translation of *SunTag-hb* mRNAs occurs in a band in nc14. (A) Anterior region of a fixed nc14 *Drosophila scFv-NG, SunTag-hb* embryo stained with DAPI (blue), anti-NG (green) and *SunTag* smFISH probes (magenta). Scale bar: 50μm. Inset regions from anterior (1) and posterior (2) regions of the expression domain. Scale bar: 5μm. (B) Quantitation of total cytoplasmic mRNAs (i), translated mRNAs (ii) and percentage of mRNAs translated (iii) per nuclear territory. Heat maps correspond to the embryo image in Fig. 3A. % EL is estimated based on an embryo length of 500μm. Nuclear territories are binned along the AP axis (bin size 20μm) and mean values per bin are determined. Mean ± SD, n = 3 embryos.

However, an alternative explanation of these data could be that the scFv-NG is limiting and has become depleted in the anterior region of the *SunTag-hb* expression domain where the mRNA number is highest in nc12-14. Therefore, we addressed this in two ways. Firstly, we introduced one and two extra copies of the maternally expressed scFv-NG fusion protein into embryos and visualised *SunTag-hb* mRNA translation at nc14. These extra copies increase the scFv-NG protein concentration in the early embryo (Fig. S2A, B). Embryos with 3 or 4 copies of the scFv-NG fusion protein still show higher translation in the posterior of the expression domain than the anterior at nc14 (Fig. S2C, D). Secondly, we removed the NLS we had added to the scFv-NG fusion protein, to increase the amount of scFv-NG protein available in the cytoplasm to bind the SunTag peptides. In the presence of maternally expressed scFv-NG (no NLS), the band of translation sites is again observed at nc14 (Fig. S2E). Together, these controls suggest that the very low proportion of *SunTag-hb* mRNAs translated in the anterior in nc14 embryos is not due to limiting scFv-NG protein, but instead reflects a repression mechanism that restricts *hb* mRNA translation to a posterior band positioned ~30-50% EL.

We also used our ability to visualise translation of *SunTag-hb* mRNAs at single mRNA resolution to address their compaction, as recent studies on a subset of mRNAs in mammalian cells have found that they shift to a more open conformation when they are being translated (Adivarahan et al., 2018; Khong and Parker, 2018). We assessed the compaction of *SunTag-hb* mRNAs during translation by using smFISH probes against the *SunTag* sequences and smiFISH probes againt *hb* sequences, positioned at the 5’ and 3’ ends of the mRNA, respectively (Fig. 4A). A representative nc14 embryo image is shown in Fig. 4Bi with two regions of interest captured at higher magnification shown in Fig. 4Bii. Using the images captured at higher magnification, we classified mRNAs as being translated or not, based on the presence or absence of a colocalised NG signal, and measured the distance between the probes following 3D rendering of the signals (Fig. 4Biii, see also Methods). This analysis reveals a range of distances between the 5’ and 3’ ends of translated and untranslated mRNAs (Fig. 4Ci shows the spread of data for a representative nc14 embryo). The median distance is significantly higher for translating mRNAs compared to those that are not, at both nc13 and nc14 (Fig. 4Cii). These data are consistent with the *SunTag-hb* mRNAs being in a slightly more open conformation during translation elongation. However, we note that the SunTag system lacks the resolution to visualise mRNAs specifically at the translation initiation step, as elongation of the ribosome through the SunTag peptides is necessary in order to classify the mRNA as being translated.

**Figure 4:**
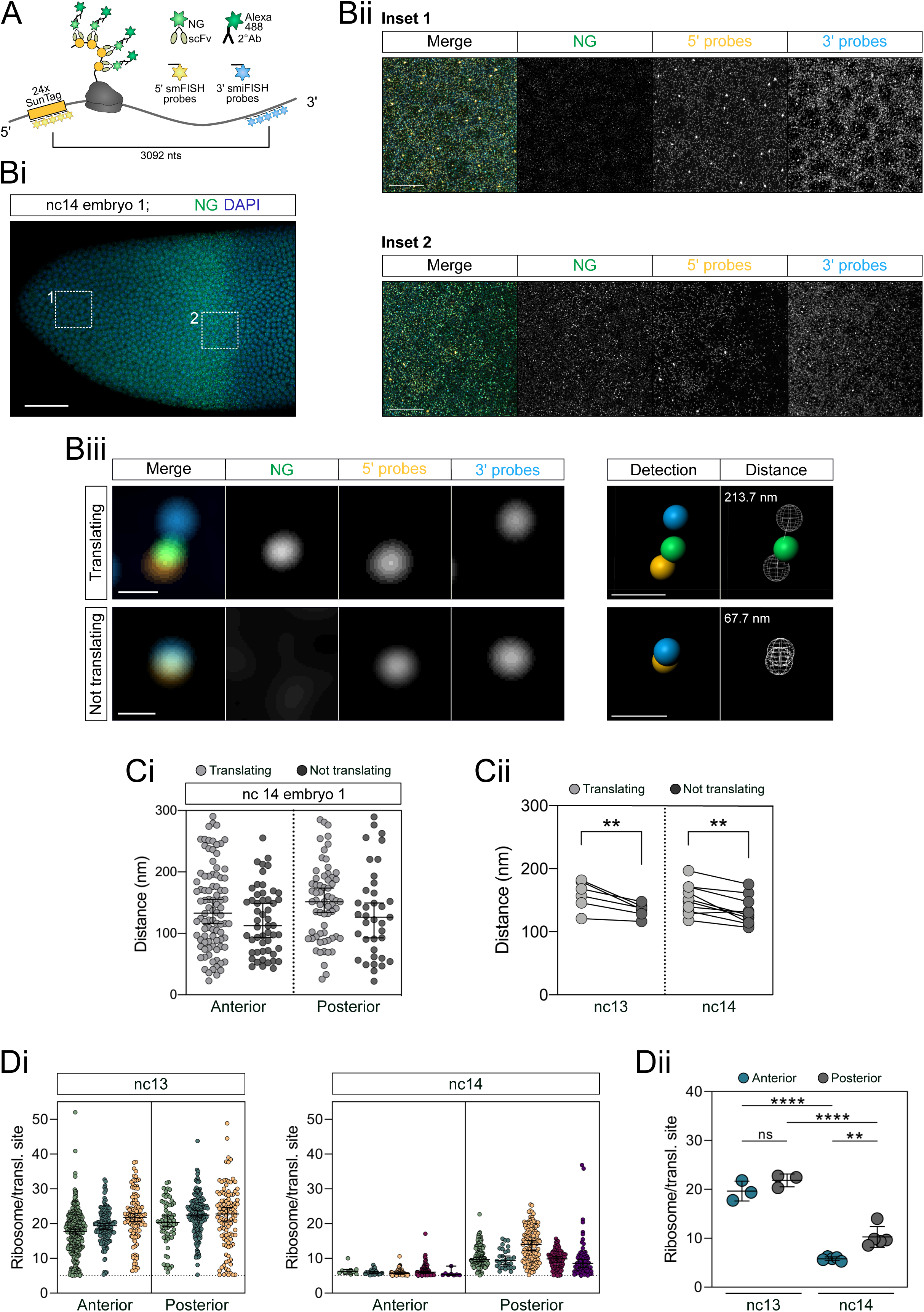
Compaction of *SunTag-hb* mRNAs and ribosome density within the expression domain. (A) Schematic showing *SunTag-hb* mRNA translation, SunTag peptides being recognised by the scFv-NG fusion protein and visualised by antibody staining. The 5’ *SunTag-hb* mRNA region is recognised by smFISH probes and the 3’UTR region by smiFISH probes, the centres of each probe set are separated by 3092 nts. (Bi) Anterior region of a fixed *scFv-NG; SunTag-hb* nc14 embryo with DAPI (blue) and anti-NG (green). (Bii) Enlarged regions from (Bi) showing an anterior (1) and posterior (2) view of the *hb* expression domain stained with *SunTag* 5’ (orange) and *hb* 3’ (cyan) probes and anti-NG (green). Here, 15 optical slices were maximum intensity projected, note that the *hb* 3’ probes also detect endogenous *hb* mRNAs. (Biii) Enlarged view of individual *hb* mRNA molecules. Translating mRNAs were identified by colocalization of the 5’ probe signal with NG. Fluorescent signals were identified using the Imaris software (Detection panel) and the closest distance between 5’ and 3’ probe signal was computed in 3D (Distance panel). Scale bars: 50 μm (Bi), 10 μm (Bii) or 0.2 μm (Biii). (Ci) Quantitation of mRNA compaction, reported as distances between the 5’ and 3’ probe sets for translating (light grey) and not translating (dark grey) mRNAs in the inset regions of Bi and (Cii) as the median of biological replicates. The lines connect the data from each biological replicate. n= 52, 92, 37 and 64 mRNA molecules (Ci) and n= 3 (nc13) and 5 (nc14) embryos (Cii). **p < 0.01, median ± 95% confidence intervals (Ci) and paired Student’s t-test (Cii). (Di) Quantitation of ribosome numbers present in translation sites in the anterior and posterior regions of nc13 and nc14 embryos. (Dii) Comparison of the average ribosome numbers between biological replicates. Ribosome numbers above 5 were detected in this analysis (see Methods for details). **p < 0.01, ****p < 0.0001; ns, not significant; median ± 95% confidence intervals (Di) and mean ± SD (Dii). n= 345, 146, 114, 65, 160, 113 translation sites (nc13) and 10, 22, 68, 99, 8, 123, 30, 138, 97, 67 translation sites (nc14) (Di). n= 3 (nc13) and 5 (nc14) embryos (Dii). One-way ANOVA with a Tukey’s multiple comparisons test (Dii).

We also used these images to estimate the number of ribosomes translating *SunTag-hb* mRNAs, based on the relative intensity of the NG signal for translation sites versus single proteins (Fig. 4D). Additionally, we applied a correction to account for the signal on the mRNA potentially arising from only partial translation of the SunTag sequence (Pichon et al., 2016) (see Methods for details). As we detect a low proportion of relatively strong NG signals that are not colocalised with mRNAs (Fig. S5D), which likely reflect scFv-NG aggregates, to classify translation sites we only quantitated NG signals that colocalised with a *SunTag-hb* mRNA signal (see Methods). We detect a range in the number of translating ribosomes on *SunTag-hb* mRNAs (Fig. 4Di), with a mean of ~20 ribosomes/mRNA in both the anterior and posterior of a nc13 embryo (Fig. 4Dii). In nc14 embryos, the *SunTag-hb* mRNAs are less heavily translated (Fig. 4Dii). Moreover, there is a spatial difference in ribosome density, with a significantly lower number of ribosomes translating *SunTag-hb* mRNAs in the anterior than posterior in nc14. Our data from nc13 and nc14 embryos suggest inter-ribosomal distances of between 180 and 690 nt on *SunTag-hb* mRNAs, which is similar to the 160-910 nt range of inter-ribosomal distances reported in mammalian cells (Morisaki and Stasevich, 2018). Together, the data presented above and our quantitation of ribosome number, suggest that in nc14 embryos not only are very few *SunTag-hb* mRNAs being translated in the anterior, but for those that are, they are only being poorly translated.

We next used the SunTag system to image translation in living embryos. Firstly, we imaged embryos that carried the *SunTag-hb* transgene and scFv-NG to visualise translation sites (Fig. 5A), in addition to Histone(His)-RFP to label the nuclei for precise ageing of the embryos. We imaged a narrow field of view in the anterior of the embryo to balance optimal temporal and spatial resolution (Movie 1, Fig. 5Bi). In this live imaging we detect a lower number of translation sites than in the fixed images, as we are only detecting the brightest translation sites. Imaging at higher magnification and with increased laser power will improve sensitivity, but the advantage of our approach here is that imaging the area extending across one third of the AP axis will provide us with an overview of the global translation dynamics. Indeed, analysis of this movie reveals how the band of translation sites in the centre of the embryo evolves during development.

**Figure 5:**
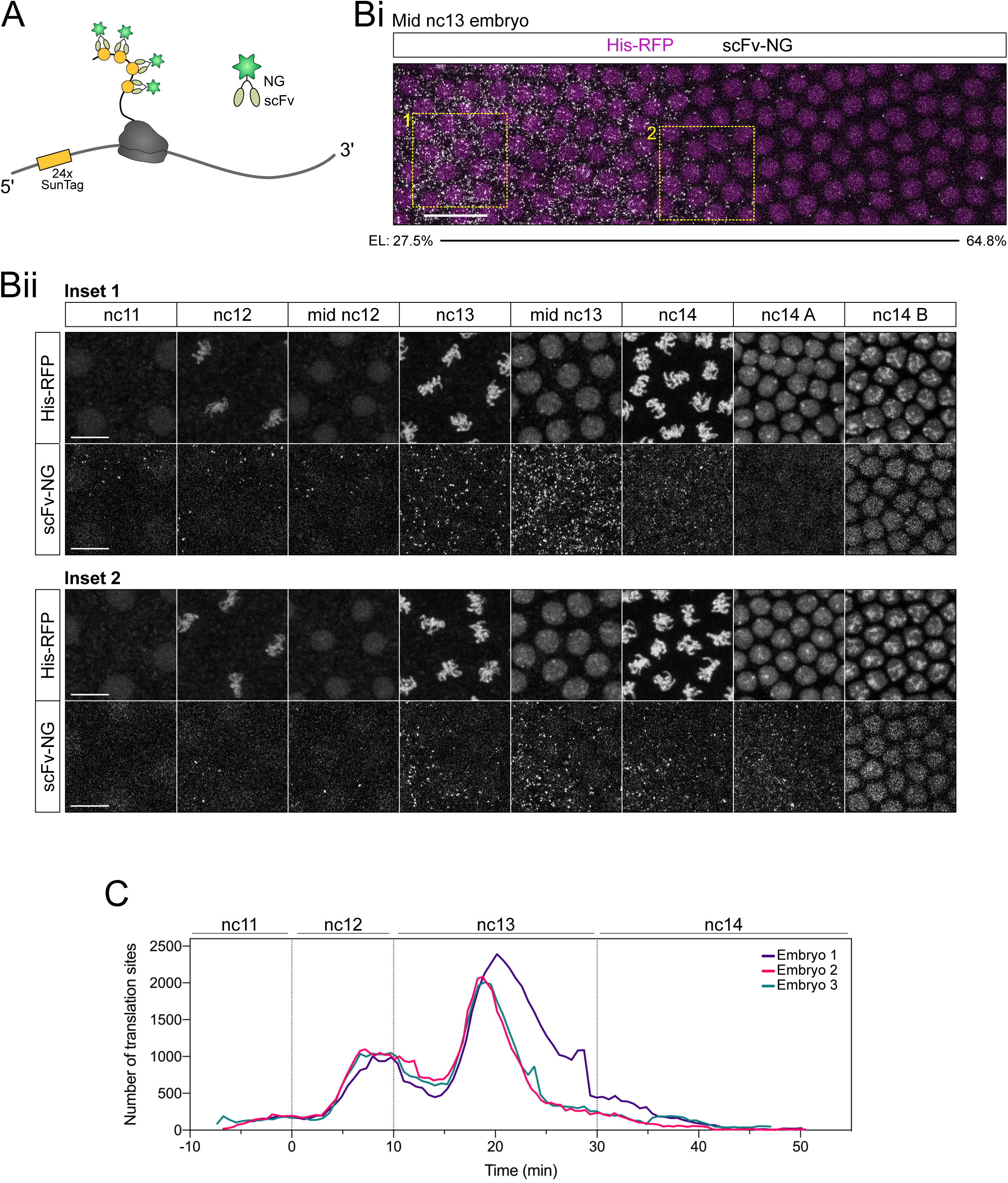
Live imaging using the SunTag system reveals global translation dynamics of *SunTag-hb* mRNAs. (A) Schematic of a *SunTag-hb* mRNA being translated and recognised by scFv-NG. (B) (Bi) Maximum projected still from Movie S1 at mid nc13 showing *hb* translation by scFv-NG fluorescence (grey) and nuclei labelled with His-RFP (magenta). (Bii) Enlarged regions from (Bi) showing the anterior (1) and posterior (2) view of *SunTag-hb* translation through developmental time. Scale bar: 20 μm (Bi), 10 μm (Bii). (C) Quantitation of translation site number over developmental time. Time-lapse datasets were aligned with t=0 min at the onset of nc12, n= 3 biological replicates.

The dynamic distribution of translation sites through developmental time is shown in stills from Movie 1, by focusing on a region in the anterior and posterior (Fig. 5Bii). These data show that, initially in nc11, translation sites are clearly visible in the anterior but not posterior regions of the expression domain (Fig. 5Bii). In nc12, translation sites are detected in both regions, although there are less in the posterior (Fig. 5Bii). In nc13 there is a large increase in the number of translation sites, again with more in the anterior, consistent with the data from fixed embryos (Fig. 2). This peak in *SunTag-hb* mRNA translation during nc13 is also obvious when the total number of translation sites is quantitated over time, for 3 biological repeats (Fig. 5C, S3B). In nc14 embryos, initially translation sites are detected in both the anterior and posterior, however over time translation sites are only detected in the posterior (Fig. 5Bii), consistent with our analysis of fixed embryos (Fig. 3). Later in nc14, no translation sites are detected and instead the intensity of the nuclear SunTag-Hb signal increases consistent with nuclear import (Fig. 5Bii).

We also used the live imaging data to plot the profiles of translation sites in bins along the AP axis over time in nc12-14, for each of the biological repeats (Fig. S3). Initially in nc12, the translation site profile is broadly similar to the mRNA distribution, with declining numbers in the posterior (Fig. S3). The number of translation sites increases throughout nc12 and in the first ~10 mins of nc13, after which the number starts to decline. Towards the end of nc13 for embryos 2 and 3, or start of nc14 for embryo 1, the translation site profiles change shape so that the position on the AP axis with the highest number of translation site moves progressively posterior (Fig. S3). There is some variation in the time at which this happens, consistent with biological variation. Nonetheless, these live imaging data show that the domain of translation contracts from the anterior over time leaving only a band of translating *SunTag-hb* mRNAs in the embryo at ~35-55% EL early in nc14, consistent with our data from fixed embryos (Fig. 3).

In order to visualise *SunTag-hb* mRNAs and translation sites in living embryos we inserted a cassette carrying 128 copies of the MS2 loops, which allows high sensitivity of detection (Tantale et al., 2016), into the 3’UTR of the *SunTag-hb* transgene (Fig. 6A). *SunTag* smFISH and NG antibody staining of fixed embryos carrying this transgene shows that insertion of the 128x MS2 cassette does not alter the mRNA translation profile (Fig. S4). Live imaging of this *SunTag-hb-MS2* transgene in embryos, which also maternally express MCP-RFP (Halstead et al., 2016) and scFv-NG, allows visualisation of the transcription site in each nucleus (Fig. 6Bi). However, the density of mRNAs in the cytoplasm and their speed of movement prevents the detection and tracking of individual mRNAs in subsequent frames (Fig. 6Bii, Movie 2).

**Figure 6:**
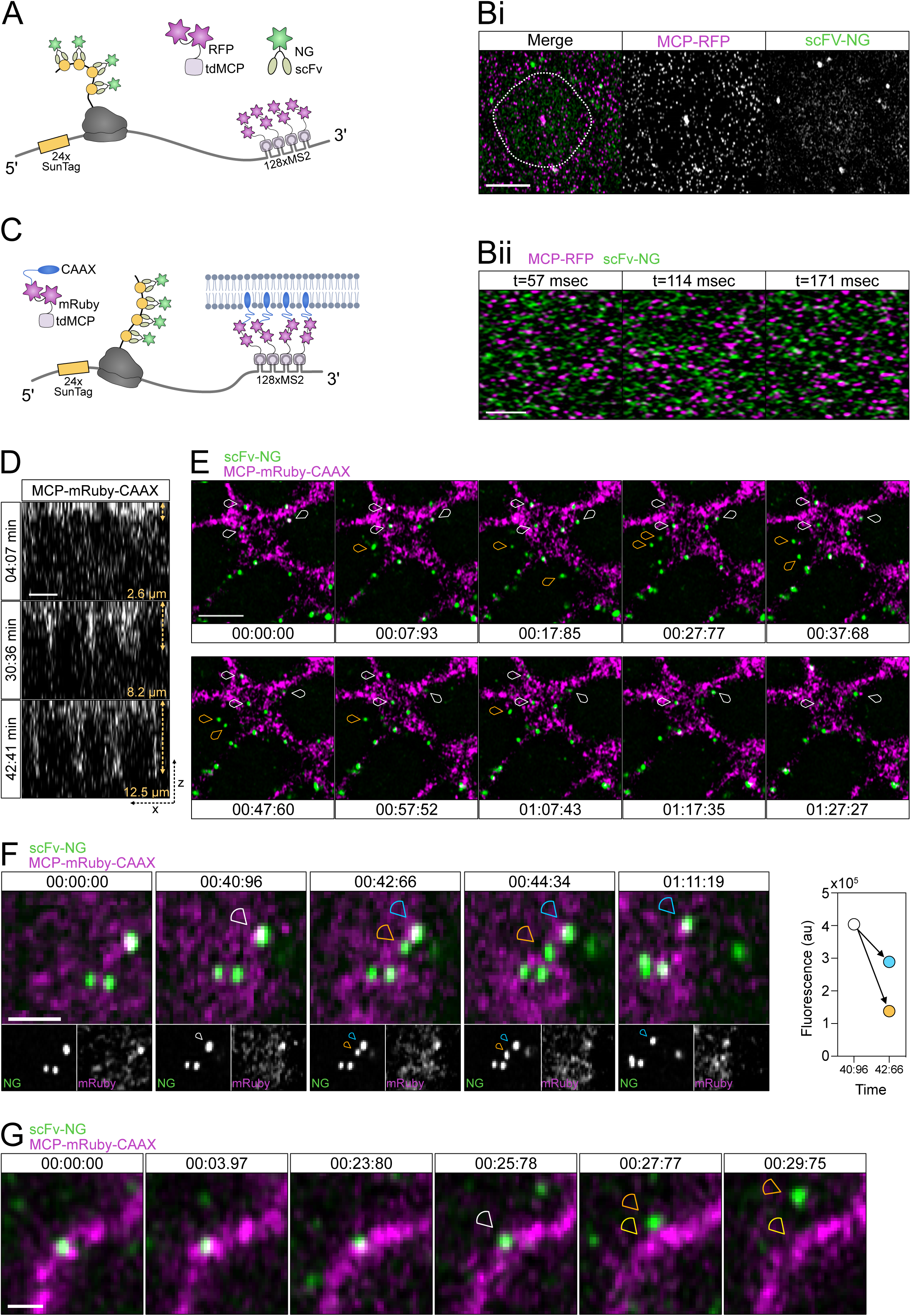
Live imaging of *SunTag-hb-MS2* mRNA translation detects different stages in the mRNA lifecycle. (A) Schematic outlining the live imaging system used to visualise both *SunTag-hb-MS2* mRNA using MCP-RFP and translation sites using scFv-NG. (Bi) A still from a live imaging movie showing that the MCP-RFP protein can be used to visualise transcription sites in the nucleus and single cytoplasmic mRNAs (magenta), in addition to visualisation of translation sites by scFv-NG proteins (green). The nuclear volume is outlined by the white, dashed line based on the nuclear scFv-NG background. (Bii) Stills from Movie S2 showing MCP-RFP and scFv-NG signals in successive frames. Scale bar: 5 μm (Bi), 2 μm (Bii). (C) Schematic showing a modified live imaging system where mRNAs are targeted to the cell membrane by the addition of a CAAX sequence to tdMCP-mRuby. Note, also tdMCP-mRuby-CAAX that is not bound to mRNA will be present in the membrane. (D) An orthoganal view of a *Drosophila* embryo during nc14 shows that MCP-mRuby-CAAX is incorporated into the plasma membrane and cellularisation can be followed by the membrane ingression. Scale bar: 5 μm. (E) Translation sites (green) tethered at the plasma membrane through MCP-mRuby-CAAX (magenta) can be followed for minutes (white arrowheads) whereas cytoplasmic proteins are visible more transiently (orange arrowheads). Time intervals are min:sec:msec, scale bar: 5 μm. See Movie S3. (F) Translation site dynamics are captured using the CAAX anchored system. A single protein (orange arrowhead, highly mobile) is detected leaving the translation site (white arrowhead), while more ribosomes continue translation of the same mRNA (blue arrowhead). Quantitation of the fluorescent signals suggest that the translation site in the plane of view contained ~4 proteins being translated at 40:96 min of which three continue being translated at 42:66 min. Time intervals are min:sec:msec, scale bar: 2 μm. See Movie S4. (G) A translation site was followed (white arrowhead) until translation was terminated, resulting in a single protein leaving the translation site (orange arrowhead, highly mobile) with the mRNA still visible (yellow arrowhead). Time intervals are min:sec:msec, scale bar: 1 μm. See Movie S5.

To facilitate longer term tracking of individual mRNAs, we used an approach described for the SunTag method in tissue culture cells, which is to add a membrane targeting CAAX sequence to MCP (Yan et al., 2016). We also expressed MCP as a tandem dimer to promote full occupancy of the MS2 stem loops and fused 2 copies of mRuby3 to MCP to avoid an associated loss of fluorescence due to MCP binding the loops as a dimer (Wu et al., 2012) (Fig. 6C). Given that the early *Drosophila* embryo is not cellularised, the tdMCP-mRuby3-CAAX fusion protein is concentrated on the available apical membrane. Consistent with this, the membrane localization of the tdMCP-mRuby3-CAAX protein allows membrane ingression to be followed in nc14 (Fig. 6D). Imaging the *SunTag-hb-MS2* transgene in embryos maternally expressing the *tdMCP-mRuby3-CAAX* transgene allows visualization of translation sites (Movie 3). Note that the embryo in Movie 3 was imaged in a single plane to allow faster temporal resolution. Stills from the movie show translation sites, that are largely immobile due to the membrane tethering, which can be tracked over time (Fig. 6E, white arrowheads). In addition, single proteins are visible that are highly mobile and move in and out of the imaged plane in successive frames (Fig. 6E, orange arrowheads). We also detect some bright NG puncta that do not colocalise with mRNAs (Fig. S5C), as described above for the fixed images. This mRNA tethering approach allows the tracking of translation sites in living embryos for >5 minutes; the NG fluorescence intensity traces of two *SunTag-hb-MS2* mRNAs in Movie 3 are shown in Fig. S5A.

This ‘long-term’ imaging of *SunTag-hb-MS2* mRNA translation sites also allows different events in the translation cycle to be visualised. For example, we have observed release of a newly synthesised SunTag-Hb protein (Fig. 6F, orange arrowhead) from the translation site (Fig. 6F, blue arrowhead, see also Movie 4). Consistent with this, the sum of the NG fluorescence intensities of the mature released protein and its translation site is equivalent to that of the translation site in the preceding frame (Fig. 6F). This quantitation also suggests that, when the protein is released, there are 3-4 ribosomes translating the mRNA in the translation site. In addition, we have been able to visualise termination of *SunTag-hb-MS2* mRNA translation (Movie 5, Fig. 6G). When the mature protein (Fig. 6G, orange arrowhead) is released from the translation site (Fig. 6G, white arrowhead), a bright mRuby3 fluorescent signal is visible (Fig. 6G, yellow arrowhead), consistent with further clustering of MCP-mRuby3-CAAX proteins when they are bound to the *SunTag-hb-MS2* mRNA. Finally, we have also visualised two translation sites that appear to repeatedly fuse then move apart (Movie 6, Fig. S5B). Whether this relates to random movement of translation sites or is associated with some regulation of translation will require further study. Together, these data show that the approaches we have described allow global translation dynamics to be elucidated, and different steps in the translation cycle to be visualised at single mRNA resolution, in living embryos in real time.

## Discussion

Here we have applied the SunTag methodology to the *Drosophila* embryo, with fixed and live imaging, to study the translation of *hb* mRNAs at single mRNA resolution during early embryogenesis. Our data from fixed embryos show uniform translation efficiency of *SunTag-hb* mRNAs across the expression domain during nc12 and nc13. We used the proximal *hb* enhancer to drive expression, which results in broadly uniform mRNA levels that decline towards the posterior border in nc12-14 (Bothma et al., 2015; Garcia et al., 2013). A similar shape of mRNA gradient is observed from the endogenous locus in nc12-13, when a shadow enhancer that activates expression in the same pattern to the proximal enhancer is also active (Bothma et al., 2015; Perry et al., 2011). Based on the profile we detect for the proportion of translated mRNAs, and the relatively constant ribosome number on mRNAs at nc13 irrespective of their position in the expression domain, our data predict that Hb protein levels decline between ~30-50% EL at nc13 (Fig 7). This is entirely consistent with a quantitative measurement of Hb levels early in nc14 (Perry et al., 2012).

**Figure 7:**
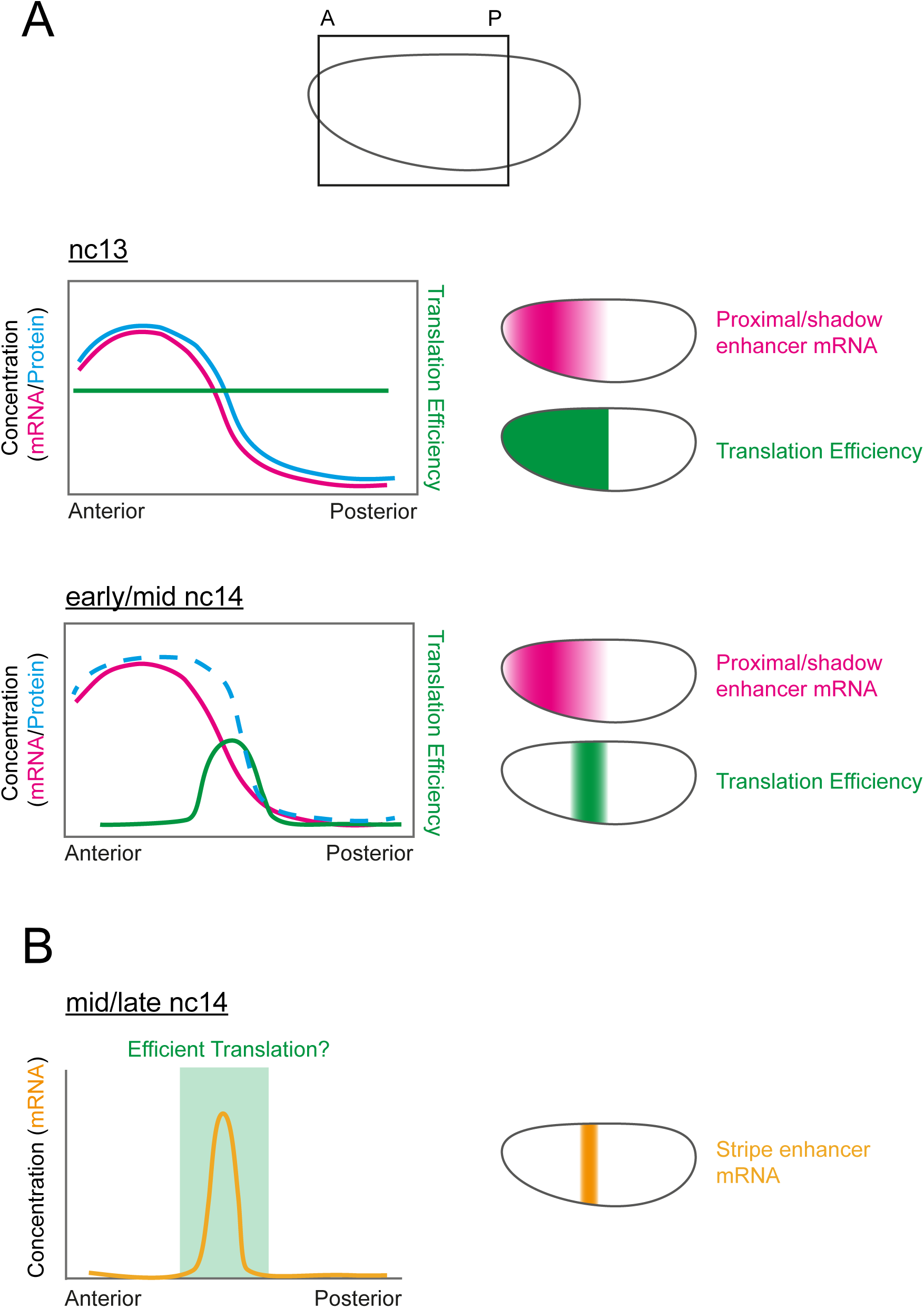
Hypothesis for the interplay between transcriptional and translational regulation of *hb*. (A) Schematic of an embryo showing the area that the graphs in the figure represent. Graphs and embryos show the approximate output from the proximal enhancer. The shadow enhancer outputs a similar pattern. In nc13, translation efficiency (green) is constant and protein concentration (blue) reflects mRNA concentration (pink). In early/mid nc14, efficient translation is limited to the posterior edge of the mRNA expression domain, sharpening the protein concentration border (dashed blue line = hypothetical output). (B) Hypothetical relationship to the stripe enhancer mRNA output. Stripe enhancer mRNA expression (yellow) occurs at the edge of the anterior expression domain. Efficient translation of this mRNA, in the area identified in (A) (nc14), would enhance the sharpness of the protein border.

In nc14, we detect less *SunTag-hb* mRNA translation overall and the distribution of translation sites changes. Analysis of fixed embryos reveals that *SunTag-hb* mRNA translation is largely repressed in the anterior but persists in a band at the posterior border of the expression domain. Our live imaging data provide a more complete picture of how the number and pattern of translation sites evolves over developmental time. There is an increase in the number of translation sites from nc11, with the highest number detected at mid-nc13. From late nc13 there is a gradual shift in the AP position of the peak of translation towards the posterior, culminating in *SunTag-hb* mRNA translation only being detected at the posterior of the expression domain in nc14 (Fig. 7). *SunTag-hb* mRNAs are still present in the anterior in nc14, but ~95% are not being translated, suggesting that they are being translationally repressed.

In addition to the highest number of *SunTag-hb* mRNA translation sites being detected at nc13, analysis of the number of ribosomes translating the mRNAs in nc13 and 14 reveals that the *SunTag-hb* mRNAs are more heavily translated in nc13. In nc14 the low number of mRNAs that are still being translated in the anterior are more poorly translated than those in the posterior, as we observe around half the number of ribosomes on anterior mRNAs in comparison to those at the posterior border. Currently the mechanism underlying repression of *SunTag-hb* mRNAs in the anterior in nc14 is unknown. As the *hb* mRNA levels are high in the region where translation repression occurs, but low where translation persists, one obvious candidate would be negative feedback by the Hb protein itself, which would take longer to reach a threshold at the posterior of the expression domain. If Hb does negatively regulate translation of its mRNA, this regulation is likely indirect as Hb lacks an obvious RNA binding domain.

The *hb* locus also has a stripe enhancer that activates transcription in a central and posterior stripe at nc14, with the central stripe positioned in the region where the early anterior expression domain ends (Margolis et al., 1995; Perry et al., 2012). As a result, the amount of Hb protein is increased in this region (~40-50% EL) in nc14 embryos, which is necessary for Hb to specify the mesothoracic T2 segment by allowing activation of *Antennapedia* (Perry et al., 2012; Wu et al., 2001). Therefore, we suggest that the translational regulation that we have visualised using the simpler system of only the proximal enhancer will also exist for the more complex pattern of *hb* mRNAs transcribed under the control of the three *hb* enhancers in the endogenous locus. Such a mechanism of prolonged translation in the posterior of the expression domain would allow *hb* mRNAs that are transcribed due to activation from the stripe enhancer to be translated, promoting development of T2. Perhaps negative feedback downstream of Hb at the translational level functions together with Hb’s ability to repress its own transcription (Little et al., 2013; Perry et al., 2012), to establish a specific level and precise pattern of Hb, which is essential for correct anterior-posterior patterning.

We propose that the increased translation efficiency of *SunTag-hb* mRNAs that we detect in the centre of the embryo at nc14, along with the extra transcription from the stripe enhancer will complement each other in nc14 to reduce heterogeneity in Hb protein expression across the expression domain and sharpen its boundary (Fig. 7) (Perry et al., 2012; Wu et al., 2001). These mechanisms may also work together to filter positional errors in the Bcd gradient, with such a noise filtering mechanism suggested for the generation of a precise Hb boundary (Houchmandzadeh et al., 2002; Yang et al., 2020). Earlier in development, in nc13, the proximal and shadow enhancers that activate transcription in the broad anterior domain function additively in the centre of the embryo, thereby maximising the transcriptional output where Bcd concentration is limiting, whereas the enhancers function subadditively more anteriorly (Bothma et al., 2015). Therefore, it appears that the embryo uses multiple mechanisms to shape the boundary of the anterior Hb expression domain during early embryogenesis. Future work studying the translation of *hb* mRNAs transcribed from the endogenous locus will allow a fuller dissection of the interplay between transcriptional and translational regulation in generating the Hb protein profiles spatiotemporally.

We also show that translated mRNAs have a tendency to be in a slightly more open conformation than untranslated mRNAs, as described in mammalian cells (Adivarahan et al., 2018; Khong and Parker, 2018). The closed loop model for translation initiation proposes an association between the 5’ and 3’ ends of the mRNA, mediated by translation factors, to explain the synergistic effects of the cap and polyA tail on translation (Vicens et al., 2018). Our findings can still be compatible with this closed loop model if, as suggested, the interaction between the 5’ and 3’ ends of the mRNA is relatively transient (Adivarahan et al., 2018; Archer et al., 2015; Khong and Parker, 2018) and/or the mRNA cycles between different conformations during the translation initiation steps (Costello et al., 2017). Alternatively, *hb* mRNA translation may be less reliant on the closed loop, with the requirement for the closed loop model for translation proposed to be mRNA dependent (Costello et al., 2015; Thompson et al., 2016). We also find no significant difference in mRNA compaction for untranslated mRNAs in the anterior and posterior in nc14 embryos, suggesting that the repression mechanism does not involve a major change in the degree of association of the 5’ and 3’ ends.

In live imaging experiments using the scFv-NG fusion and standard MCP-fluorescent protein fusion, we found that the speed with which mRNAs move in 3D and the density of mRNAs in the cytoplasm precludes tracking of individual mRNAs over time. Two strategies have been used to overcome this same problem in mammalian cells, including tethering the mRNAs to the membrane (Yan et al., 2016) or studying local protein synthesis on the endoplasmic reticulum (Wu et al., 2016). By exploiting the membrane tethering approach (Yan et al., 2016), we show that translation of individual mRNAs in the embryo can be monitored for at least 6 minutes, allowing events such as release of nascent proteins or translation termination to be captured.

The different MCP and scFv lines that we have generated will facilitate the quantitative study of translation of other *Drosophila* mRNAs, in live and/or fixed samples. Consistent with this, we note that another study, carried out in parallel to ours, has also recently described the application of the SunTag approach to the study of *twi* mRNA translation in the *Drosophila* embryo (Dufourt et al., 2020). Overall, the quantitative data from the fixed SunTag embryo images and dynamic information about translation sites gained from imaging live embryos represents a powerful combination for probing the spatiotemporal regulation of translation. Future studies can also exploit the many advantages of the early embryo, such as the synchronous nuclear divisions that will allow the regulation of translation during mitosis to be studied without the need for drugs to arrest the cell cycle. Moreover, the SunTag approach can be used to reveal, at unparalleled spatiotemporal resolution, how the integration of transcription and translation dynamics shapes developmental patterning.

## Materials and Methods

### Generation of plasmids and fly lines

#### phbP2>24xGCN4-hb-24xMS2, phbP2>24xGCN4-hb-128xMS2 and phbP2>24xGCN4-hb

The *hb* P2 enhancer and 5’UTR, coding sequence and 3’UTR were PCR amplified from genomic DNA. The SunTag (24 x GCN4 peptides) was PCR amplified from the plasmid pcDNA4TO-24xGCN4_v4_sfGFP (Addgene #61058) (Tanenbaum et al., 2014), the 24xMS2 stem loop sequence was purified from pCR-24xMS2L-stable (Addgene_31865) by restriction digest and the 128xMS2 from pMK123-128xMS2 (Tantale et al., 2016). The *hb* P2 enhancer, 5’UTR, 24xGCN4 and *hb* CDS were inserted into the StuI/BamHI sites of pUASp (DGRC #1358) using multiple-insert In-Fusion cloning. The *hb* 3’UTR was inserted into the BamHI/NdeI sites. 24xMS2 and 128xMS2 were inserted into the BamHI site. The sequences of all primers used are shown in Supplemental Table 1. The attP/attB system was used for specific integration of the transgenes into site 86Fb (chr3) by the University of Cambridge microinjection service.

#### pCasper-nos>noNLS-2xMCP-2xRuby3-CAAX-tub3’UTR+1kb

2xMCP and mRuby3 were cloned into pCasper4 containing an attB site, the *tubulin* 3’UTR+1kb downstream in the XbaI site and the *nos* promoter in the EcoRI site. The NLS 2x MCP from pHsp83-NLS-HA-2xMCP-2xTagRFP-T (Addgene 71242) (Halstead et al., 2016)was inserted into KpnI/SpeI. 2xmRuby3 with a short linker was generated by introducing a second copy of mRuby3 into the BglII site of pKanCMV-mRuby3-10aa-H2B (Addgene 74258). The two copies of mRuby3 were then subcloned into the SpeI site of pCasper-nos>NLS-HA-2xMCP-tub3’UTR+1kb. The CAAX from pHR-PP7-2xmCherry-CAAX (Addgene 74925) (Yan et al., 2016) was inserted between SpeI/BamHI. The NLS was subsequently removed by digest with KpnI/NheI, the overhangs generated were filled using Klenow Fragment and the plasmid was re-ligated. Transgenes were injected into w1118 using random P-element insertion. A line with the transgene located on the 3^rd^ chromosome was used for this study.

#### pCasper-nos>ScFv-mNeonGreen-GB1-NLS-tub3’UTR+1kb

The scFv sequence was obtained from plasmid pHR-scFv-GCN4-sfGFP-GB1-NLS-dWPRE (Addgene 60906) (Tanenbaum et al., 2014). scFv-mNeonGreen-GB1-NLS was assembled using In-Fusion^®^ HD Cloning Plus multiple insert cloning (Takara Bio #638911) and integrated into the KpnI/SpeI sites of pCasper-attB-nos>tubulin3’UTR+1kb (containing the *nos* promoter in EcoRI to direct maternal expression and *tubulin* 3’UTR+1kb of downstream sequence in XbaI). To generate the noNLS version the scFv-mNeonGreen-GB1 was amplified by PCR and inserted into KpnI/NotI of pCasperattB-nos>tub3’UTR. The attP/attB system was used for specific reintegration of the transgene into sites 25C6 (chr2) and 86Fb (chr3) by the University of Cambridge microinjection service.

### Fly stocks

Fly stocks used are: *y^1^w^67c23^* (used as wildtype, BL6599), His-RFP (BL23650), nosP>scFv-mNeonGreen-GB1-NLS, nosP>scFv-mNeonGreen-GB1, nosP>scFv-mNeonGreen-GB1-NLS; scFv-mNeonGreen-GB1-NLS/MKRS, nosP>scFv-mNeonGreen-GB1-NLS; scFv-mNeonGreen-GB1-NLS, hbP2>24xGCN4-hb, hbP2>24xGCN4-hb-128xMS2, nosP>tdMCP-2xmRuby3-CAAX and w-; if/CyO; P{w+, pHsp83-NLS-HA-2xMCP-2xtagRFP}9/TM3 Ser (Halstead et al., 2016). Flies were maintained at 18C and crosses performed at 25C. Flies were raised on standard fly food (yeast 50g/L, glucose 78g/L, maize flour 72g/L, agar 8g/L, nipagen 27ml/L, and propionic acid 3ml/L). For smFISH/IF experiments, embryos were laid on apple juice agar plates with yeast paste for three to four hours at 25°C prior to fixation.

### Embryo Fixation

Embryos were dechorionated for 2 minutes in 50% bleach and washed with water/Triton NaCl solution. Embryos were transferred to 1:3:4 37% formaldehyde: heptane: fixing solution (1.33x PBS, 0.07M EGTA (pH8.0)) and shaken for 20 minutes at 300 rpm. Embryos were devitellinised by the addition of methanol and shaking/vortexing.

### smFISH/smiFISH and Immunofluorescence

Fixed embryos were rocked at room temperature in glass scintillation vials in a series of washes – 5 minutes 50:50 methanol:PBT (1x phosphate buffered saline + 0.1% Tween-20); 4 x 10 minutes PBT; 10 minutes 50:50 PBT:Stellaris wash buffer (10% formamide, 2x SSC); 2 x 5 minutes Stellaris wash buffer. Embryos were then allowed to settle for 5 minutes in Stellaris hybridization buffer (100mg/ml dextran sulfate, 10% formamide, 2x SSC). Stellaris hybridization buffer was removed and replaced with fresh hybridization buffer, and embryos were incubated in a 37°C water bath for 2 hours as a pre-hybridization step. Probes sets for smFISH (Biosearch Technologies) or smiFISH (Sigma Aldrich) were diluted in Stellaris hybridization buffer to final concentration 100nM (smFISH probes) and 80nM (smiFISH probes) and embryos were incubated with probes in the dark at 37°C for at least 14 hours. Following probe hybridization, embryos were kept in the dark. Embryos were washed for 30 minutes, rinsed and washed for 3 x 15 minutes at 37°C, all with Stellaris hybridization buffer pre-warmed to 37°C. Embryos were then washed for 15 minutes at room temperature in Stellaris wash buffer, followed by washing 3 x 10 minutes in PBT. Embryos were blocked for 30 minutes in PBT + 1x WBR (Western Blocking Reagent, Sigma, #11921673001) and then incubated overnight at 4°C with primary antibody (mouse anti-mNeonGreen [32F6], Chromotek, 1:250) diluted in PBT + 1x WBR. Embryos were rinsed twice in PBT, washed 4 x 15 minutes in PBT and blocked for 30 minutes in PBT + 1x WBR. Embryos were incubated with secondary antibody (donkey anti-mouse Alexa Fluor 488, Thermo Fisher Scientific, 1:250) diluted in PBT + 1x WBR for 2 hours at room temperature. Embryos were rinsed twice and washed 4 x 15 minutes in PBT, with the third wash including DAPI at 1:1000. Embryos were mounted on slides in ProLong Diamond Antifade Mountant (Thermo Fisher Scientific, Cat# P36961). smFISH/smiFISH probe sequences are shown in Supplemental Table 1.

### Static image acquisition

Images used for the analysis of mRNA and translated mRNA number were acquired using a Leica TCS SP8 AOBS inverted microscope using a 63x/1.40 oil objective and 0.75x confocal zoom. The confocal settings were pinhole 1 airy unit, scan speed 400Hz unidirectional line scanning, 2x line averaging and format of 2048 x 2048 pixels. Images were collected sequentially with either Photon Multiplying Tube Detectors or Hybrid Detectors and illuminated using a white laser at 70%. The following detection mirror settings were used: Photon Multiplying Tube Detector DAPI excitation at 405nm (2%, collection: 415-470nm); Hybrid Detectors: AlexaFluor 488 excitation at 490nm (10%, collection: 500-540nm) and Quasar 570 excitation at 548nm (15%, collection: 558-640nm). Optical z-stacks were acquired at 300nm spacing.

Images used for compaction analysis and quantitation of ribosome number were acquired with a Leica TCS SP8 AOBS inverted microscope using a 100x/1.40 HC PL Apo CS2 oil objective and 3x confocal zoom. The confocal settings were as follows, pinhole 0.65 airy unit, scan speed 400Hz bidirectional line scanning, 4x line averaging and a format of 4096 x 4096 pixels. Images were collected sequentially with either Photon Multiplying Tube Detectors or Hybrid Detectors and illuminated using a white laser at 70%. The following detection mirror settings were used: Photon Multiplying Tube Detector DAPI excitation at 405nm (11%, collection: 415-470nm); Hybrid Detectors: AlexaFluor 488 excitation at 490nm (12%, 1 to 6us gating, collection: 500-540nm), Quasar 570 excitation at 548nm (20%, 1 to 6us gating, collection: 558-640nm) and Quasar 670 excitation at 647nm (21%, 1 to 6us gating, collection: 668-779nm). Optical z-stacks were acquired at 200nm spacing.

Images of embryos with different number of maternal scFv-NG copies were acquired with a Leica TCS SP8 AOBS inverted microscope using a 40x/1.30 HC PL Apo CS2 oil objective and 1x confocal zoom. The confocal settings were as follows, pinhole 0.8 airy unit, scan speed 400Hz bidirectional line scanning, 4x line accumulation and a format of 4752 x 4752 pixels. Images were collected sequentially using the same laser and detector settings as described above.

All raw images were deconvolved using Huygens Professional software (SVI) and figures show maximum intensity projections unless stated otherwise in the figure legends. All embryos are oriented with the anterior to the left. In the figure panels that focus on the anterior region of the embryo, the image is rotated and shown on a black background.

### Live imaging microscopy

Embryos were dechorionated in bleach and mounted onto a Lumox imaging dish (Sarstedt AG & Co, Cat#94.6077.305) as described (Hoppe and Ashe). To visualise translation sites globally together with maternally deposited His-RFP, embryos were imaged using an LSM 880 Airy Upright microscope with an Airyscan Fast detector and an EC Plan-Neofluar 40x/1.30 oil DIC M27 objective at 1.1x optical zoom. 25 planes were acquired with 850nm spacing at 2996 x 788 pixels, with a z-stack acquisition time of ~40s. His-RFP was excited by the 561nm laser line at 1.5% laser power and scFv-NG by the 488nm laser line at 5%.

Live translation and the MCP-RFP bound *SunTag-hb* RNA were imaged using an LSM 980 Airy Upright microscope with an Airyscan Fast detector to facilitate fast sampling times. Images were acquired using a PL Apo 63x/1.40 Oil objective at 8bit, 17x optical zoom, with 2x line averaging and bidirectional scanning at an LSM scan speed of 9. MCP-RFP was excited by the 594 nm laser line at 7% laser power and scFv-NG by the 488 nm laser line at 2.2% laser power. A single plane time series was acquired at 176×160 pixel and with a frame time of ~57 msec.

All other time-lapse datasets showing live translation were acquired on the LSM 880 Airy Upright Airyscan Fast setup using a PL Apo 63x/1.40 Oil objective, unidirectional scanning at 16bit and fluorophore excitation using the 594 nm laser line at 15% laser power and 488 nm laser line at 15% laser power. Additional specifications were as follows:

MCP-RFP and scFV-NG: Images showing nascent hb transcription were acquired at 8x optical zoom, 252×252 pixel, 2 optical slices 400 μm spaced apart and a time resolution of ~842 ms per frame. A single plane is shown in in Figure 6Bi.

MCP-Ruby-CAAX and scFV-NG: Single plane time-lapse datasets were acquired at 8x optical zoom and with 184×184 pixel (overview and translation termination) and a time resolution of ~2 sec per frame or at 116×116 pixels (single protein release) with a time resolution of ~1.7 sec per frame.

MCP-Ruby-CAAX orthogonal view: An embryo was imaged throughout nc14 at 0.7x optical zoom, 2176×1384 pixel. An optical stack was acquired with 1 μm spacing and a total depth of 16 μm, resulting in a final time resolution of ~123 sec.

### Western blot

*Drosophila* embryonic extracts were prepared based on a published protocol (Guilgur et al., 2014; Prudêncio and Guilgur, 2015). 1-4 hour old embryos were dechorionated and homogenised in ice cold lysis buffer (150mM NaCl, 20mM Tris.HCl (pH8), 2mM EDTA, 0.1% NP-40, 1mM DTT, 1x cOmplete™, EDTA-free Protease Inhibitor Cocktail Sigma #11873580001). Extracts were centrifuged at 15,000xg for 20 minutes at 4°C. Lysates were denatured in NuPAGE™ LDS Sample Buffer (4X) (Thermo Fisher Scientific, #NP0007) with 50mM DTT. A standard Western blot protocol was used with anti-HA(12CA5) (1:1000, Merck #11583816001), anti-Actin(20-33) (1:1000, Merck #A5060), anti-mNeonGreen [32F6], (1:1000, Chromotek) primary antibodies and IRDye^®^ 680RD Donkey anti-Rabbit IgG (H+L) (1:10,000, Li-Cor #926-68073), IRDye 800CW Donkey anti-Mouse IgG (H+L) (1:10,000, Li-Cor #926-32212) secondary antibodies. Signals were detected using Li-Cor Odyssey^®^ CLx Infrared detection system.

### Quantitation of the number of total and translated mRNAs

For analysis of mRNA and translation site number, fixed embryos were stained with SunTag smFISH probes, anti-mNeonGreen antibody and DAPI, and imaged using the acquisition details described above. *SunTag* mRNAs and mNeonGreen foci were identified using the Imaris software (Imaris software 9.2; Bitplane, Oxford Instruments, Concord MA) ‘spots’ function. *SunTag* mRNA puncta were identified with spot volume 0.6μm in diameter and 1.2μm in the z direction. mNeonGreen foci were identified with spot volume 0.5μm in diameter and 1μm in z. Nuclei were identified using the Imaris ‘surfaces’ function. Translated mRNAs were identified using Imaris ‘Co-localise Spots’ Xtension, with spots called as colocalised if the distance between the centroids of the two spots was less than 0.6μm. Spots inside the nucleus were identified using Imaris ‘Split Into Surface Objects’ Xtension. Nuclear spots were excluded by combining data for all spots and removing any duplicated spots. Spot IDs were then used to extract statistics for cytoplasmic spots, which were used for analysis.

Cytoplasmic spots were assigned to surfaces using a modified version of spotMe.py from (Hoppe et al., 2020), spotMe_EmbryoMid.py. Spot assignment to surfaces remains unchanged, while midline detection is modified to fit the midline of the embryo rather than the expression domain. As only a portion of the embryo is obtained in each frame, and the orientation of the embryo can vary, the midline is detected by identifying the edge nuclei of the embryo. Nuclei within 1.5 internuclear distances of the edge of the frame are removed. Centroids for all permutations of unique nuclei pairs from opposing edges of the embryo are calculated creating a coordinate set representing the mean position between the two embryo edges. These mean positions are used to calculate the midline using a least-squares approach. The anterior position of the embryo is determined by identifying the end of the embryo with the least nuclei in bins along the x-axis. Positions along the AP axis of the embryo for each nucleus are determined by assigning nuclei to their closest position on the AP axis then calculating the distance between that AP position and the position closest to the anterior-most nuclei. Code used can be found at https://github.com/TMinchington/sass. In nc14 embryos with either the *SunTag-hb* or *SunTag-hb-MS2* transgenes, we observed 25/26 embryos with a band of translation sites and 1/26 embryo with uniform translation sites. As we only found a single nc14 embryo with uniform translation sites, we consider that it had a developmental defect. 3 embryos with a band were fully analysed.

### Compaction and ribosome number

For compaction and ribosome number analysis fixed *Drosophila* embryos were stained with smFISH probes against the *SunTag* coding sequence, smiFISH probes against the *hb* coding sequence, and anti-mNeonGreen antibody. The two probe sets, targeting different regions of the *hb* transgene, are spaced 3092 nts between their centres. Anterior and posterior regions were imaged with a 100x objective and 3x optical zoom (acquisition details described above). Regions of these images were analysed for compaction and ribosome number quantitation. Compaction analysis: *SunTag* (5’ probe) and *hb* (3’ probe) mRNA foci as well as mNeonGreen translation sites were identified using the Imaris “spots” function (Imaris software 9.2; Bitplane, Oxford Instruments, Concord MA) and their statistics exported. The closest distance in three dimensions between a *SunTag* spot and its corresponding *hb* spot was computed based on the spot centroids using a custom python script. Foci were assumed to be from the same mRNA if they were less than 300 nm apart. mRNAs were classified as translating if the distance between a *SunTag* spot centroid and the nearest mNeonGreen centroid was less than 300nm.

Ribosome number analysis: To quantify the number of ribosomes within a translation site, the fluorescence intensity of *SunTag* mRNA and mNeonGreen signal was quantified using the AirLocalize software (Trcek et al., 2017). AirLocalize outputs were used to perform a colocalization analysis using custom python scripts to classify the mNeonGreen signal into translation sites and single proteins. The number of ribosomes present in translation sites, which were assigned based on colocalization with a mRNA, was calculated using the median fluorescence intensity of single proteins and corrected for the position of the GCN4 repeats. A correction factor was used, calculated by (length_SunTagCDS_ × 0.5 + length_hbCDS_)/length_total_, to account for ribosomes that have only translated a fraction of the SunTag repeat (Pichon et al., 2016). Ribosome numbers larger than five could be confidently identified and are presented in Figure 4Di-ii.

### Nuclear scFv-NG concentration

Nuclear scFv-NG levels were quantitated from static images of *Drosophila* embryos whose mothers carried 2x, 3x or 4x copies of scFv-NG in their genome. Nuclei positioned outside of the *hb* expression domain, in the posterior region of the embryo, were segmented using the “surface” function in Imaris. The sum fluorescence intensity of a nucleus in the scFv-NG channel was divided by the nuclear area. Background levels were estimated using “spot” objects placed in the cytoplasm between posterior nuclei. The sum fluorescent background value was divided by the spot area and subtracted from the nuclear fluorescence.

### Nuclear tracking and translation site identification in live imaging datasets

Nuclei were segmented and tracked in the Imaris software based on the His-RFP fluorescent signal. Nuclei were segmented using the “surface function” individually for each nuclear cycle and then tracked through time using the inbuilt autoregressive motion tracking with a maximum frame gap size of 5 and a maximum travel distance of 6 μm. The scFv-NG channel was first smoothed and blurred using a wavelet filter (Imaris X-tension; Hoppe and Ashe, 2021) and then translation sites were identified using the Imaris “spots” function. Translation sites were estimated to 0.7 μm across with a z-axis point spread function estimation of 1.4 μm. To determine the background fluorescence in the translation site channel, a set of “spots” of the same volume was generated for background correction. Statistics were exported for tracked surfaces, translation site spots and background spots. Translation sites were linked to their closest nucleus (nuclear territories) and background correction was applied using the custom python script ‘sass’ (https://github.com/TMinchington/sass) (Hoppe et al., 2020).

### Translation site fluorescence and tracks

Fluorescent translation sites were quantified using the Imaris “spots” function and sum fluorescence values are reported in Fig. 6F. Spot sizes were chosen to contain the full translation site fluorescence. Background fluorescence was determined using spots of the same size as translation spots and the sum fluorescence was subtracted from translation sites. To track translation site fluorescence through time, the Imaris inbuilt spot tracking function was used. Individual fluorescent traces of translation sites through time are shown in Figure S5.

### Statistical Analysis

Statistical comparisons were performed using one-way ANOVA or Kruskal-Wallis-Tests with multiple comparison, two-tailed Student’s t test or paired Student’s t tests using GraphPad Prism and R (Version 3.5.2). The statistical test used and the respective sample size can be found in the figure legends. Statistical significance was assumed at p<0.05 and individual p values are indicated in the figure legends.

## Acknowledgments

We thank Mark Ashe for helpful discussions, Nathan Garnham for help generating the transgenes, Edouard Bertrand for the pMK123-128xMS2 plasmid, Frank Wippich and Anne Ephrussi for the MCP-RFP flies, Shane Herbert for microscopy time, and the University of Manchester Bioimaging Facility for support. This work was funded by a Wellcome Trust Investigator Award to H.L.A. (204832/Z/16/Z) and Wellcome Trust PhD studentship to D.V. (215206/Z/19/Z).

## Author Contributions

Conceptualization: D.V., H.L.A.; Investigation: D.V., C.H., C.S.; Software, T.G.M; Writing: D.V., H.L.A.; Supervision: H.L.A.; Funding Acquisition: H.L.A.

