## Supplementary Material for "Dynamics of *hunchback* translation in real time and at single mRNA resolution in the *Drosophila* embryo"

**Figure S1: Variation of mRNA numbers within embryos.**

(A) Quantitation of mean number of total *SunTag-hb* mRNAs (i), translated *SunTag-hb* mRNAs (ii), and percentage translated (iii) per nuclear territory for three nc12 biological repeat embryos. Mean ± SD, n = 3.

(B) The number of total (i) and translated *SunTag-hb* mRNAs (ii) for each individual nuclear territory is plotted across the AP axis from one nc12 embryo.

(C, D) As in (A, B) for nc13 embryos.

(E, F) As in (A, B) for nc14 embryos.

**Figure S2: Visualisation of *SunTag-hb-128* mRNAs in embryos with altered scFv-NG levels or subcellular localisation.**

(A) Western blot with anti-NG and anti-HA to detect the scFv-NG protein (the scFv-NG fusion protein has a HA tag at its N-terminus, see Fig. 1A) and actin as a loading control. Extracts are from 1-3 hr old embryos that are wildtype (0x) or from females with 2 or 4 copies of the *scFv-NG* transgene. A representative blot is shown, the graph shows quantitation of the HA signal relative to actin, the mean ± S.D from 3 repeats is plotted on the graph. ∗∗p < 0.01, two-tailed Student's t test.

(B) Quantitation of nuclear scFv-NG fluorescence in posterior nuclei outside of the *hb* expression domain, in embryos from females with the indicated number of copies of the *scFv-NG* trangene. ∗∗∗∗p < 0.0001, median ± interquartile range and Kruskal-Wallis-Test with Dunn's multiple comparisons test. n= 160, 209, 129 nuclei in embryos from females expressing 2, 3 and 4 copies of *scFv-NG*, respectively.

(C, D) Anterior region of fixed nc14 *Drosophila* embryos, from females carrying 3 (C) or 4 (D) copies of the *scFv-NG* transgene and stained with smFISH probes against the *SunTag* region (magenta), anti-NG (green) and DAPI (blue). A translation band is visible at the edge of the *SunTag-hb* expression domain. Scale bars: 50 µm.

(E) As in (C) but the embryo is from a female expressing *the scFv-NG-noNLS* transgene and the *SunTag-hb* transgene contains 24xMS2 in the 3'UTR.

**Figure S3: Average translation site number across the AP axis from live imaging data.**

**(A)** Mean number of *SunTag-hb* translation sites along the AP axis over time. Translation sites are grouped into positional bins along the embryo's AP axis and time points are split according to their nuclear cycle. Bin width = 20 µm, time resolution is indicated in the graphs.

(B) Average number of translation sites detected per time point compared between different nuclear cycles. ∗∗p < 0.01, ∗∗∗∗p < 0.0001. Mean ± SD and one-way ANOVA with a Tukey's multiple comparisons test, n= 3 biological replicates.

**Figure S4: Visualisation of translation of *SunTag-hb-MS2* mRNAs.**

Anterior views of nc12, 13, 14 embryos from females maternally expressing the scFv-NG transgene crossed to males carrying the *SunTag-hb-MS2* transgene (with 128xMS2), stained with DAPI (blue), anti-mNeonGreen (green), and SunTag smFISH probes (magenta). Scale bar: 50μm.

**Figure S5: Live imaging translation traces and scFv-NG aggregates.**

(A) Still from a live imaging movie showing *SunTag-hb-MS2* mRNA translation sites (green) in embryos expressing MCP-mRuby-CAAX (magenta) and scFv-NG proteins. Fluorescence intensity traces of two translation sites (arrowheads) are shown for more than 6 min of developmental time. See Figure 6E for stills and Movie S3. Scale bar: 5 μm.

(B) Still from a movie as in (A), scale bar: 5 μm. Inset shows separate translation sites (white arrow heads) that fuse (orange arrowhead) and separate throughout time, scale bar: 2 μm. Time intervals are min:sec:msec. See Movie S6 for inset and Movie S3 for original time-lapse dataset.

(C) Orthogonal view of time-lapse live imaging data showing the occurrence of aggregates (orange arrowheads) where bright scFv-NG fluorescence (green) does not overlap with single mRNA molecules (magenta). Scale bar: 5 μm.

(D) Orthogonal view of a static image from a fixed *Drosophila* embryo showing translation sites (white arrowheads) and an aggregate (orange arrowhead), where the NG (green) signal does not overlap with the *SunTag-hb* mRNA signal (magenta). Our quantitation of the number of mRNAs being translated in fixed embryos is based on colocalization of the NG signals with mRNAs, therefore these aggregates are not counted as translation sites (see Methods).

**<u>Movies:</u>**

**Movie S1:** Maximum intensity projection of a representative embryo showing *hb* translation sites (grey) and Histone-RFP (red) through nc11-14. Embryos are from females with single copies of *scFv-NG* and *His-RFP* crossed to males with the *SunTag-hb* transgene. Imaged with a 40x objective, 1.1 optical zoom and 46 sec time resolution between frames. Time stamp is in min:sec.

**Movie S2:** Single plane time-lapse experiment visualising cytoplasmic *SunTag-hb* mRNA bound by MCP-RFP (magenta), and translation sites or mature proteins bound by scFv-NG (green). Embryos are from females carrying single copies of the *scFv-NG* and *MCP-RFP* transgenes crossed to *SunTag-hb-MS2* transgenic males. Imaged with a 63x objective and a time resolution of ~ 57 msec per frame. Time stamp is in sec:msec.

**Movie S3:** Single plane time-lapse experiment visualising plasma membrane tethered MCP-mRuby-CAAX and bound *SunTag-hb* mRNAs (magenta) and translation sites or mature proteins bound by scFv-NG (green). Embryos are from females carrying single copies of the *scFv-NG* and *tdMCP-Ruby-CAAX* transgenes crossed to *SunTag-hb-MS2* transgenic males. Imaged with a 63x objective, 8x optical zoom, and a time resolution of ~ 2 sec per frame. Time stamp is in min:sec.

**Movie S4:** Single plane time-lapse experiment showing a single protein being released from a translation site (green, scFv-NG) and diffusing away from the mRNA (magenta, MCP-mRuby-CAAX) and site of translation. The embryo genotype and imaging settings are as described for movie 3, except the time resolution is ~ 1.7 sec per frame.

**Movie S5:** Single plane time-lapse experiment visualising translation termination. An active translation site site (green, scFv-NG) terminates translation and a single protein diffuses away from the mRNA molecule (magenta, MCP-mRuby-CAAX). The embryo genotype and imaging settings are as for Movie S3.

**Movie S6:** Enlarged view of Movie S3, highlighting independent translation sites (green) fusing and separating over time.

Vinter_Figure S1

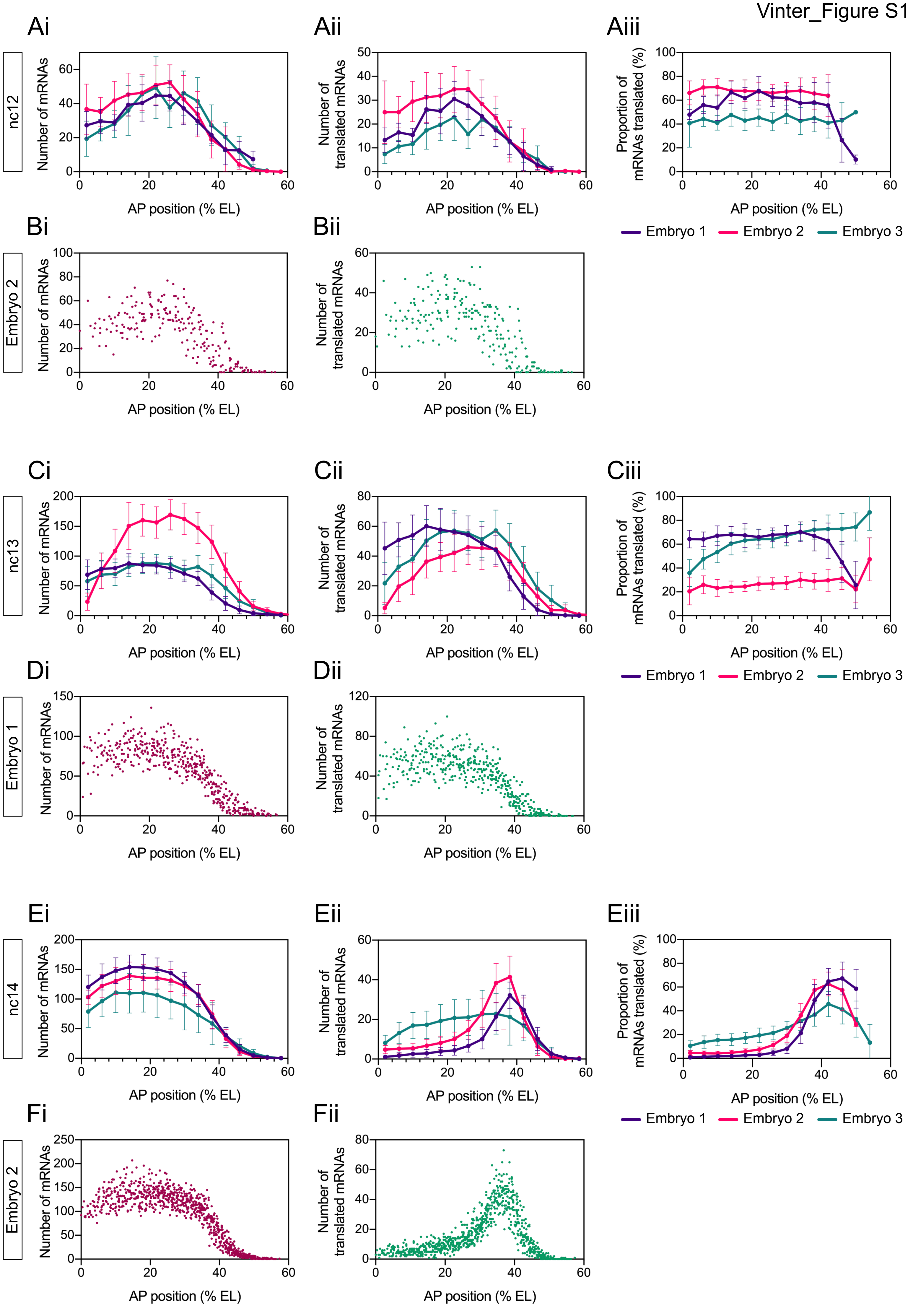

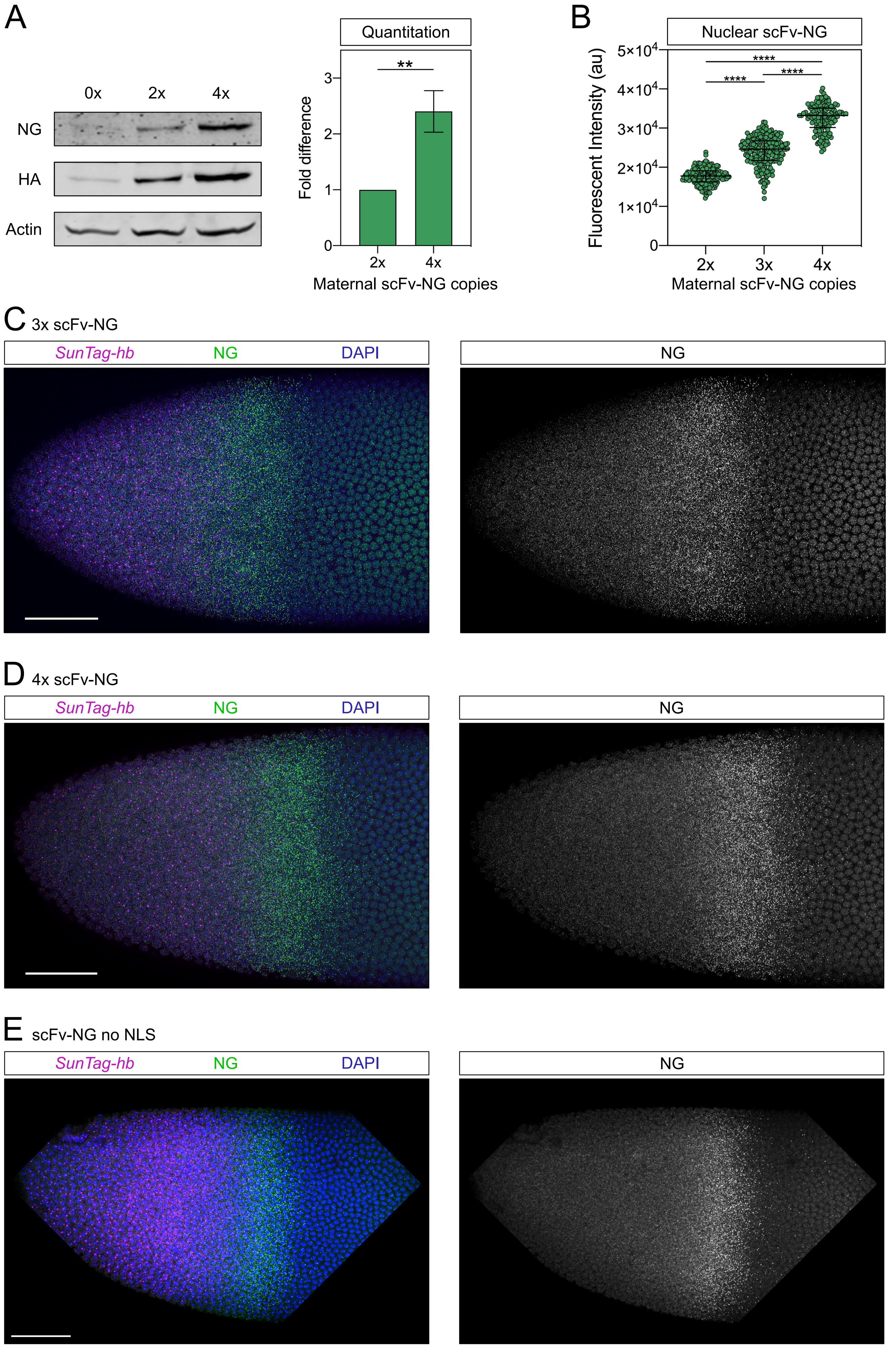
A
0x
2x
4x
NG
HA
Actin
Quantitation
**
Fold difference
0
1
2
3
2x
4x
Maternal scFv-NG copies
B
Nuclear scFv-NG
****
****
****
Fluorescent Intensity (au)
$5\times10^4$
$4\times10^4$
$3\times10^4$
$2\times10^4$
$1\times10^4$
0
2x
3x
4x
Maternal scFv-NG copies
C 3x scFv-NG
SunTag-hb
NG
DAPI
NG
D 4x scFv-NG
SunTag-hb
NG
DAPI
NG
E scFv-NG no NLS
SunTag-hb
NG
DAPI
NG

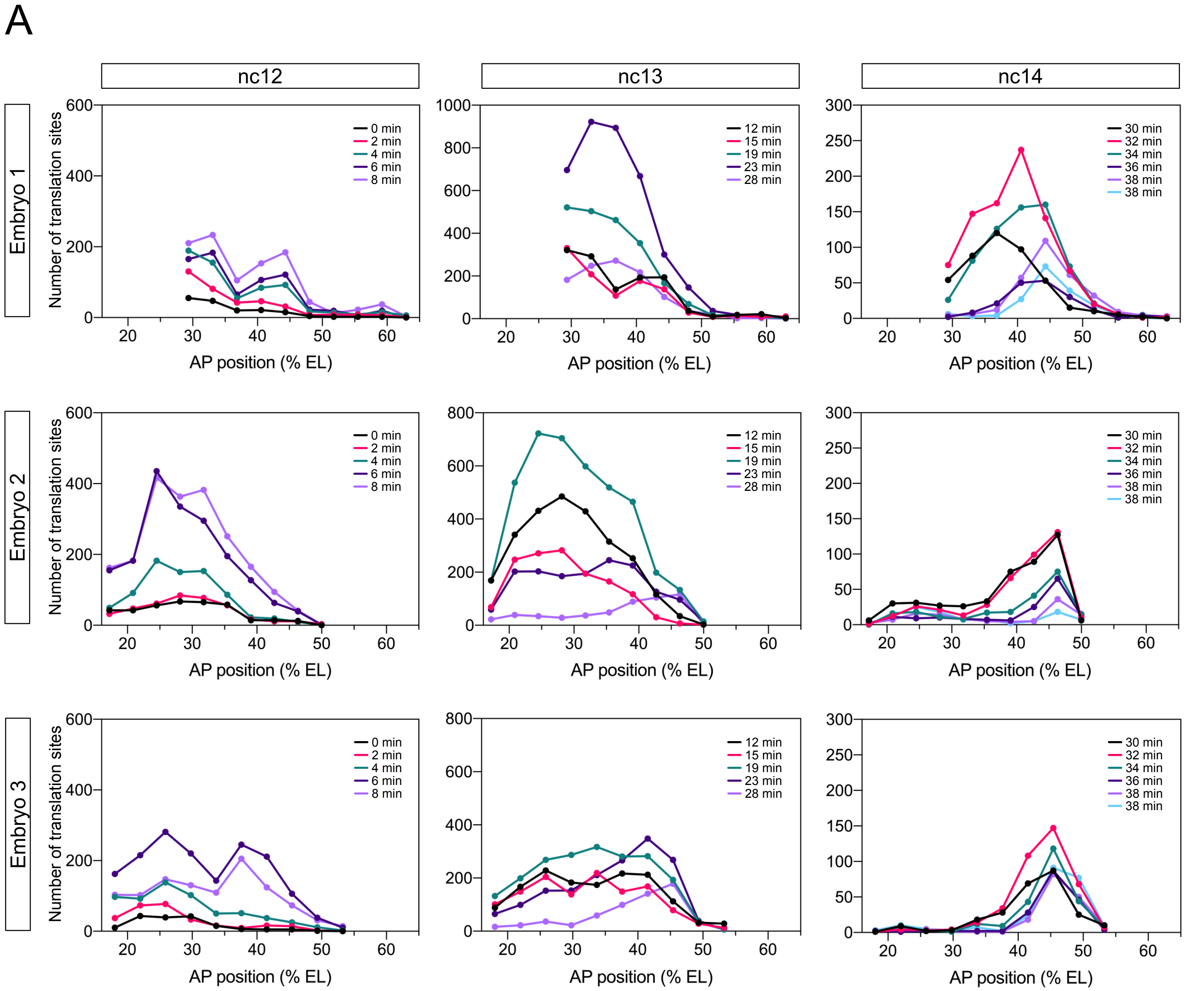
A
nc12
nc13
nc14
Embryo 1
Embryo 2
Embryo 3
Number of translation sites
AP position (% EL)
0 min
2 min
4 min
6 min
8 min
12 min
15 min
19 min
23 min
28 min
30 min
32 min
34 min
36 min
38 min
38 min

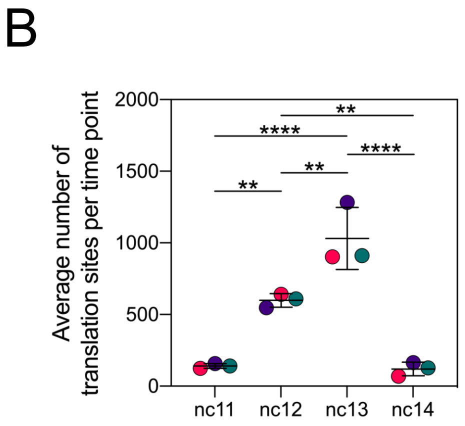
B
Average number of translation sites per time point
nc11
nc12
nc13
nc14
**
****
**
****
**

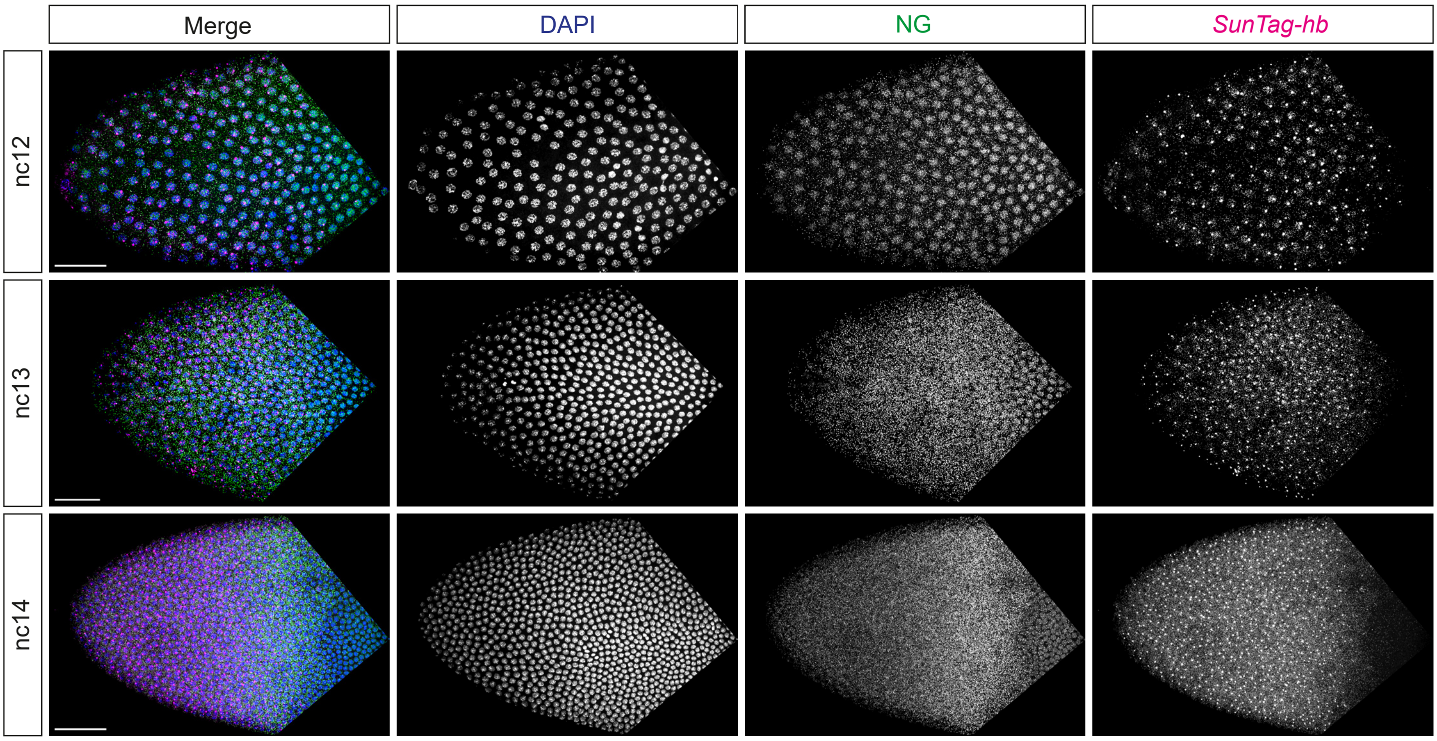
Merge
DAPI
NG
SunTag-hb
nc12
nc13
nc14

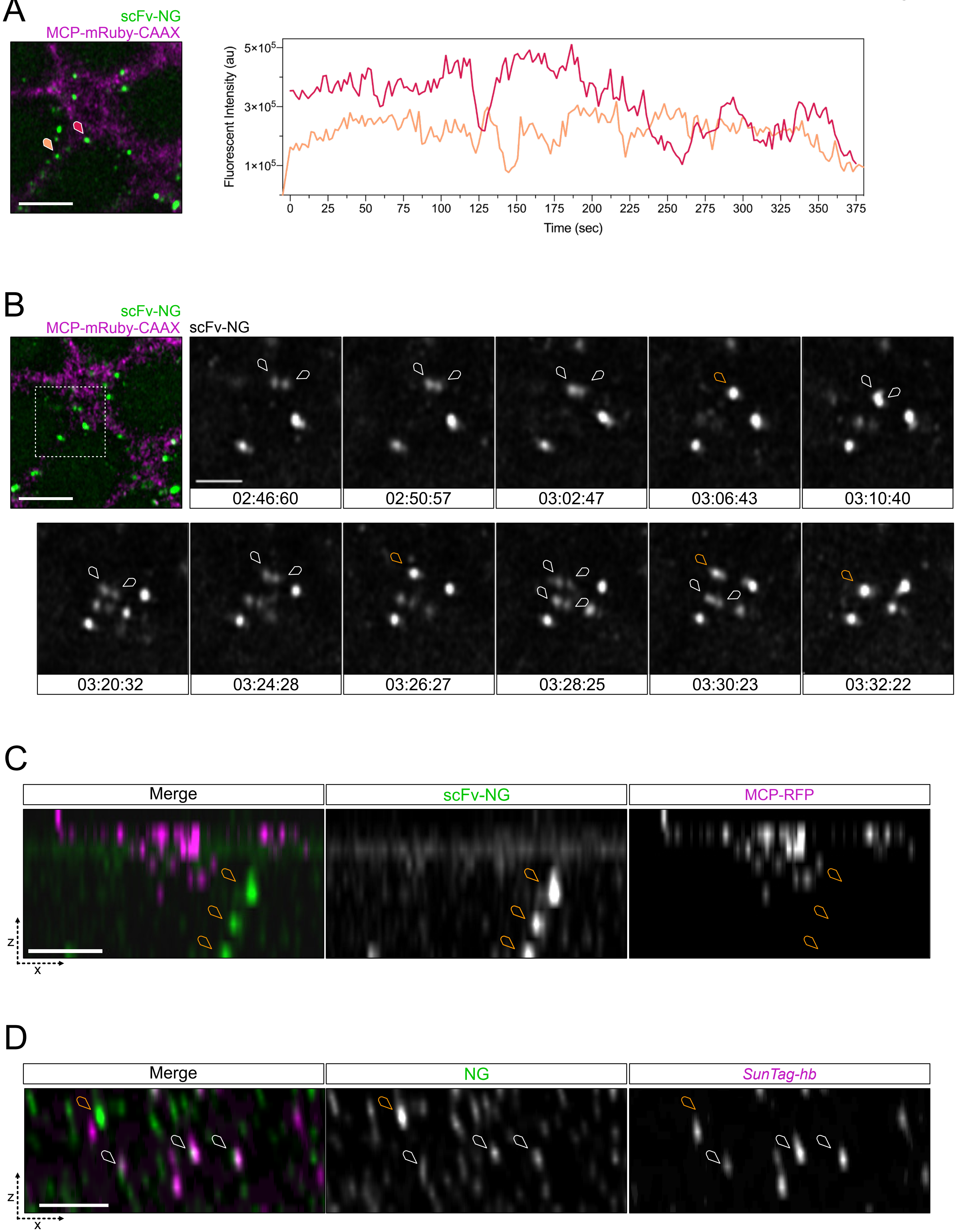
A
scFv-NG
MCP-mRuby-CAAX
Fluorescent Intensity (au)
5×10^5
3×10^5
1×10^5
0
25
50
75
100
125
150
175
200
225
250
275
300
325
350
375
Time (sec)
B
scFv-NG
MCP-mRuby-CAAX
scFv-NG
02:46:60
02:50:57
03:02:47
03:06:43
03:10:40
03:20:32
03:24:28
03:26:27
03:28:25
03:30:23
03:32:22
C
Merge
scFv-NG
MCP-RFP
z
x
D
Merge
NG
SunTag-hb
z
x

**Supplemental Table 1: Primers and Probe sequences.**

| | |
|---|---|
| **Primers** | |
| hb 5'UTR StuI F | tcgaggtcatcaagcttaggccttaactacgaaactgcccacgc |
| hb 5'UTR R | gttcttccatcttggcggctctagacggc |
| GCN4 F | agccgccaagatggaagaacttttgagcaaga |
| GCN4 R | cccagttctgctttttaagtcgggctacttca |
| hb CDS F | acttaaaaagcagaactgggagacgacagcc |
| hb CDS BamHI R | gaggtcgactctagaggatccttaggagtgagcattcctggcc |
| hb 3'UTR BamHI F | tcactcctaaggatccgttccccatcaccatcacc |
| hb 3'UTR NdeI R | tccaaagcacgcatatgatgaaagctaaagtcatagtcagg |
| NLS_MCP KpnI F: | gggtaccgcaatggcccaaaaaag |
| NLS_MCP SpeI R: | aaactagtactctagaatccgcgtagatgc |
| CAAX SpeI F | actagtacatatgggtggaggttctggt |
| CAAX BamHI R | gggatccggccgccatatgttacatg |
| Ruby3 1 NheI F | aaccgtcagatccgctagc |
| Ruby3 1 SpeI R | aagactagtcttgtacagctcgtccatg |
| Ruby3 2 BglII R | tagatctgagtccggacttgtac |
| Ruby3 2 BglII F | catagatctatggtgtctaagggcgaag |
| ScFv F | gacatacgaattcggtacctcacgcgtgccaccat |
| ScFv R | tcgcccttgctcaccataccgcttccacctccac |
| mNeonGreen F | gtggaggtggaagcggtatggtgagcaagggcga |
| mNeonGreen R | ctcttcggtccgacctcccttgtacagctcgtccatgc |
| GB1-NLS F | gcatggacgagctgtacaagggaggtcggaccgaagag |
| GB1-NLS R | tctagaggatccagatccactagagctgccttgtaagtcattgg |
| GB1-noNLS R | gcggccgcttattcggttaccgtgaaggt |
| **SunTag smFISH probes** | |
| SunTag_1 | ccacttcgttctcaagatga |
| SunTag_2 | gccagaacctttcttaagac |
| SunTag_3 | ttgaaagcagttcttctcca |
| SunTag_4 | ctcattttccaggtggtaat |
| SunTag_5 | aatttttgctcagcaactcc |
| SunTag_6 | ttctttagtcgtgctacttc |
| SunTag_7 | tttcgagagtaactcctcac |
| SunTag_8 | ccacttcgttttcgagatga |
| SunTag_9 | gataatagctcttctccaga |
| SunTag_10 | tcattttcgaggtggtagtt |
| SunTag_11 | acttccctttttaagcgtg |
| SunTag_12 | tcttggatagtagctcttca |
| SunTag_13 | ctacctcgttctcaagatga |
| SunTag_14 | tagttcttcgagagcagttc |
| SunTag_15 | gatccctttttaatcgagc |
| SunTag_16 | tgaaagtagttcctcaccac |
| SunTag_17 | cttcgttttcgaggtggtaa |
| SunTag_18 | ccctgaacctttctttaatc |
| SunTag_19 | tactcagtaattcttcaccc |
| SunTag_20 | ctacctcattttccagatga |
| SunTag_21 | tttcgatagcaactcttcgc |
| SunTag_22 | tttttgagcctagcaacttc |
| SunTag_23 | agtggtagtttttcgagagc |
| SunTag_24 | tttgctcaataactcctcgc |
| SunTag_25 | cgcgacttcgttctctaaat |
| SunTag_26 | ttcgataagagttcttcgcc |

| | |
|---|---|
| SunTag_27 | ctcattttcgaggtggtagt |
| SunTag_28 | gtggtagttcttgctcaag |
| SunTag_29 | attcttgctgagcaattcct |
| SunTag_30 | cgacttcgttctccaaatga |
| SunTag_31 | tttactcaacaattcctccc |
| SunTag_32 | cgacttcattttccaagtgg |
| SunTag_33 | ttgctcaataactcttcgcc |
| SunTag_34 | ttcgttctccaagtggtaat |
| SunTag_35 | agttcttcgataagagctcc |
| SunTag_36 | gcgacttcattctctaagtg |
| SunTag_37 | agtggtagttcttgctcaag |
| SunTag_38 | ttagatagtaactcttcccc |
| SunTag_39 | cctcgttctcgagatgataa |
| SunTag_40 | gatagttcttcgacaggagt |
| SunTag_41 | cctttttaagtcttgcaacc |
| SunTag_42 | ttcttactgagtagttcctc |
| SunTag_43 | tcctgatcctttcttcaaac |
| SunTag_44 | cttttgagagcagttcttcg |
| SunTag_45 | gcaacctcattttccaaatg |
| SunTag_46 | tgccacttcccttttttaaa |
| SunTag_47 | tttcgacagaagttcctcac |
| SunTag_48 | taagtcgggctacttcattc |
| **hb 3' smiFISH probes** | |
| hb3'_1 | CCAGCTTCTAGCATCCATGCCCTATAAGctccaccttgagattcatgg |
| hb3'_2 | CCAGCTTCTAGCATCCATGCCCTATAAGattcgagctgctcatcagag |
| hb3'_3 | CCAGCTTCTAGCATCCATGCCCTATAAGttgaggacgcgtcccttg |
| hb3'_4 | CCAGCTTCTAGCATCCATGCCCTATAAGgttgtaacagggtgtcca |
| hb3'_5 | CCAGCTTCTAGCATCCATGCCCTATAAGatgtcatggcctccgatc |
| hb3'_6 | CCAGCTTCTAGCATCCATGCCCTATAAGactttcagttgctcggga |
| hb3'_7 | CCAGCTTCTAGCATCCATGCCCTATAAGagttggcatgggtgtgct |
| hb3'_8 | CCAGCTTCTAGCATCCATGCCCTATAAGcggcaatgggcgaggatg |
| hb3'_9 | CCAGCTTCTAGCATCCATGCCCTATAAGtcctcgggcatgggtttg |
| hb3'_10 | CCAGCTTCTAGCATCCATGCCCTATAAGgaactggtgcccgagcag |
| hb3'_11 | CCAGCTTCTAGCATCCATGCCCTATAAGcgtctccatcgactcatc |
| hb3'_12 | CCAGCTTCTAGCATCCATGCCCTATAAGcactggtattggcctgcg |
| hb3'_13 | CCAGCTTCTAGCATCCATGCCCTATAAGctggacgccgtcgaactg |
| hb3'_14 | CCAGCTTCTAGCATCCATGCCCTATAAGtattgctgctggcattggag |
| hb3'_15 | CCAGCTTCTAGCATCCATGCCCTATAAGttccattgctgctggaattg |
| hb3'_16 | CCAGCTTCTAGCATCCATGCCCTATAAGcttgcactcgtagatggc |
| hb3'_17 | CCAGCTTCTAGCATCCATGCCCTATAAGcagtacttgcactcgtagat |
| hb3'_18 | CCAGCTTCTAGCATCCATGCCCTATAAGggcgtccttgaagaagatat |
| hb3'_19 | CCAGCTTCTAGCATCCATGCCCTATAAGccatgtgaatggtgtagagc |
| hb3'_20 | CCAGCTTCTAGCATCCATGCCCTATAAGcacatcgtcgcagctgtg |
| hb3'_21 | CCAGCTTCTAGCATCCATGCCCTATAAGccgcacatgttgcacttg |
| hb3'_22 | CCAGCTTCTAGCATCCATGCCCTATAAGcgaagaggccgacgggtc |
| hb3'_23 | CCAGCTTCTAGCATCCATGCCCTATAAGgcattcctggccatgtga |
| hb3'_24 | CCAGCTTCTAGCATCCATGCCCTATAAGaacggatccttaggagtg |
| hb3'_25 | CCAGCTTCTAGCATCCATGCCCTATAAGcaaggtgatggtgatggg |
| hb3'_26 | CCAGCTTCTAGCATCCATGCCCTATAAGttctggacaacgattatatgat |
| hb3'_27 | CCAGCTTCTAGCATCCATGCCCTATAAGgtttggaaaacttatgctacga |
| hb3'_28 | CCAGCTTCTAGCATCCATGCCCTATAAGggcttatgtacaattttcga |
| hb3'_29 | CCAGCTTCTAGCATCCATGCCCTATAAGtaggtctagaattagcggct |
| hb3'_30 | CCAGCTTCTAGCATCCATGCCCTATAAGggtggctacagttcaataca |

| | |
|---|---|
| hb3'_31 | CCAGCTTCTAGCATCCATGCCCTATAAGgtgtataggagacagattga |
| hb3'_32 | CCAGCTTCTAGCATCCATGCCCTATAAGgttttcagggtttttagtcg |
| hb3'_33 | CCAGCTTCTAGCATCCATGCCCTATAAGtccatgcatttatgatagtttt |
| hb3'_34 | CCAGCTTCTAGCATCCATGCCCTATAAGgatttaacttaggcttatgttt |
| hb3'_35 | CCAGCTTCTAGCATCCATGCCCTATAAGgttgtttcgcttgactcaac |
| hb3'_36 | CCAGCTTCTAGCATCCATGCCCTATAAGtttgactttggactgttggt |
| hb3'_37 | CCAGCTTCTAGCATCCATGCCCTATAAGacatactcattatgtatatgtt |
| hb3'_38 | CCAGCTTCTAGCATCCATGCCCTATAAGgcgcatatacgaatatgttagg |
| hb3'_39 | CCAGCTTCTAGCATCCATGCCCTATAAGatcagagaaagaactgagtgtt |
| hb3'_40 | CCAGCTTCTAGCATCCATGCCCTATAAGggcgaattcgctttcaacta |
| **Flap sequences** | |
| Z_flap | cttatagggcatggatgctagaagctgg |
